# Effect of quantified cranial osteopathic manipulation on wild type and transgenic rat models of Alzheimer’s disease

**DOI:** 10.1101/2024.11.05.621645

**Authors:** De’Yana Hines, Hope Tobey, Patrick Dugan, Seth Boehringer, Richard Helm, Ramu Anandakrishnan, Stephen Werre, Pamela VandeVord, Blaise M. Costa

## Abstract

Alzheimer’s Disease is a chronic progressive neurodegenerative disorder that impairs the cerebral lymphatic system and compartmental fluid exchange leading to a decline in cognitive function. Due to the lack of disease modifying medications, non-pharmacological Cranial Osteopathic Manipulation (COM) is evolving as a potential minimally invasive treatment choice. In this work, the effect of quantified COM treatment, using a nanosensor glove, on 3-month-old (Yg) and 18-month-old transgenic (Tg) rat model of Alzheimer’s Disease were studied using the Morris Water Maze (MWM), Western Blots, and Proteomics and Transcriptomics assays. The results revealed that COM had minimal to no significant difference in the behavioral and biochemical parameters in the Yg rats, suggesting COM treatment was harmless. While COM exhibited no significant differences in Tg rat MWM escape latency, navigation to the platform was significantly different on testing days 5 and 6, with p-values of signed initial heading error were 0.014 and 0.034 respectively. This indicates a difference in learning and spatial working memory. A proteomic assay on Tg rat hippocampus identified 51 significantly differentially expressed proteins with 34 associated with neurological disorders, while transcriptome remained indifferent. In this study, for the first time we have established a technique to quantify the force applied during COM treatment on an animal model of AD, offering a more objective approach for evaluating the effect of such treatments. Our results indicate that a quantifiable COM can be applied to rodents and to study the resulting behavioral and biochemical phenotypes.

## Introduction

Alzheimer’s Disease (AD) is a chronic progressive neurodegenerative disorder that causes cognitive impairment leading to severe memory deficits over time. AD is in the top 10 leading causes of death in the US(1–3), sixth in 2019 and seventh in 2020 and 2021, affecting over 6 million Americans(4). Current treatments for AD primarily inhibit cholinesterase enzyme (ex. Donepezil) or NMDA subtype of glutamate regulators (ex. memantine)(5), rather than modifying pathogenic mechanisms such as reduction in cerebral vasculature pulsation and impaired cerebral fluid circulation(6,7). The FDA has recently approved amyloid specific monoclonal antibodies such as lecanemab which target the pathophysiology of AD. However, amyloid related imaging abnormalities (ARIA) are known associated side effects that can pose severe neurological problems(8). This presents a need for non-pharmacological methods with minimal to no adverse effects to treat the underlying mechanisms and symptoms of AD. Based on the principle of the Cranial Osteopathic Manipulation (COM), which is believed to improve the cerebral fluid circulation and change the expression of proteins involved in the metabolic waste clearance(9,10), there is a potential for COM to meet the clinical need.

COM is a clinically practiced treatment that requires minimal invasive intervention in the form of physical contact. The mechanistic principle behind the treatment is similar to the lymphatic pump treatment, which is another osteopathic manipulative technique that is more commonly used in clinics(11). Use of COM to alter the function of the glymphatic and meningeal lymphatic pathways, which could improve fluid flow and metabolic waste clearance, evolved based on recent findings associated with clearance performed by the central nervous system’s (CNS) lymphatic vessels(12–15). A study showed that COM altered amyloid beta (Aβ) levels via astrocyte activation, upregulating aquaporin and lymphatic vessels, and modulated synaptic transmission through excitatory and inhibitory pathways in an AD rat model(9). Spatial memory was also reported to be improved with COM treated rats as they exhibited a shorter escape latency time compared to their untreated counterparts of the same age. Studies investigating other models of neurological injuries and disorders report significant differences in treatment groups post-COM (16,17). COM treatment was found to increase interstitial fluid flow along with an improvement in anxiety-like behaviors in traumatic brain injury (TBI) preclinical models (16). These findings strengthen the idea that COM could be a clinical strategy to mechanically improve cerebral fluid circulation. Another study investigating COM as a method for migraine-like pain mitigation, found that COM blocked the development of cutaneous allodynia suggesting use as a preventative or abortive treatment(17). Collectively, these results cannot be directly compared between the studies due to a difference in animal disease models. However, the influences of COM treatment in improving relevant symptoms of neurological disorders remain consistent throughout the studies. This necessitates further research into the mechanistic benefits that quantifiable COM has as a treatment to mitigate the development of AD.

In the present study, we have employed a nanosensor glove to quantify the force applied on both sexes of wild type and Tg 344 rats and assessed the spatial and biochemical outcomes of the treatment.

## Methods

### Animals

Animal experiments and housing has been approved by the Institutional Animal Care and Use Committee (IACUC) of Virginia Tech (Protocol ID# 15-099 and ID#19-045). Two cohorts of rats were used for this study. (1) Twelve 3-month-old wild type male and female rats (Yg) were purchased from Charles River laboratories and (2), eleven 18-month-old Tg F344 male and female rats (Tg) were purchased from Rat Resource and Research Center (RRRC) at the University of Missouri. Their cohort expressing two Swedish point mutations in the Amyloid Precursor Protein encoding gene (K595N & M596L) and exon 9 deletion in the presenilin 1 gene. Both cohorts of rats were randomly divided into two groups, untreated (UT) and COM treated (COM). All rats were provided with normal food and water ad libitum. A 12-hour light-dark cycle was chosen for their housing arrangements. All the methods described were conducted according to relevant guidelines and regulations.

### COM Treatment

COM treatment was provided every day for seven days by a board-certified osteopathic physician. Rats were anesthetized with 1.5-3% isoflurane throughout the COM procedure. Untreated rats were also anesthetized for the average duration of treatment to nullify the influence from isoflurane in behavioral testing. During COM treatment, the osteopath physician applied mechanical pressure over the rat’s occipital squama which puts tension around the 4^th^ ventricle. The slight pressure applied intends to improve symmetry of the cranial rhythmic impulse (CRI), impact fluctuation of the CSF-ISF fluid compartment, and improve cranial bone and dural membrane mobility. Completion of treatment is identified once the operator feels the tissue relax. Successful completion ideally improves symmetry and or fullness of the CRI. Treated rats received 2-4 ½ minutes treatment each session. FingerTPS (Medical Tactile, Inc.) nanosensor gloves, were used to record real-time pressure and duration of treatment for quantification purposes. Clinical notes were taken to document the specifics of COM treatment, CRI changes, and any additional observable changes.

### Morris Water Maze Assay

The Morris Water Maze (MWM) assay was used to study spatial learning and memory. MWM was performed every day for 8 days. A combination of latent and repeated learning approaches was taken. Latent learning requires the animal to be placed on the platform before each trial while repeated learning requires a set of reversal or serial phase shifts(18,19). The shifts in this assay are differing starting positions for each testing day. A combination of both learning paradigms helps identify spatial awareness and the animal’s flexibility to learn across different shifts.

On day 0, the rats were introduced to the MWM tub (5ft diameter, 2ft high walls) for acclimatization. Using AnyMaze (AnyMaze 5.1, Stoelting Co. IL), a video tracking software, the tub was divided digitally into four quadrants. A platform zone (∼5in diameter) was digitally overlayed on the platform located in the northwest quadrant. Water was kept to room temperature and water level was roughly 1-2cm below the platform for learning days. Visual cues, to serve as landmarks, of a triangle, circle, and square were attached to the edge of the maze in the northwest (NW), northeast (NE), and southeast (SE) quadrants zones. On learning trials (days 1-4), the rats were trained every day in each of the four quadrants [NW, NE, SE, and southwest (SW)] to find the platform. For these days the water was transparent, and the platform was visible to the rats. Rats were given 60s to reach the platform after leaving, with their face towards the wall, into one of the four quadrants. Before each trial the rats were placed on the platform for 15s. Rats that could not reach the platform within 60s were gently guided to the platform. After finishing the trials rats were gently wiped dry and placed under a heating lamp for two minutes. On testing days (5-8), the water was made opaque, and the water level was raised to make the platform hidden. On day 8 the platform was removed. Two probe trials were performed for each animal from two different starting locations (example, Trial 1:SE zone, Trial 2: NW zone. These trials are performed for three days (day 5,6&7) and one for day 8, with the same exposure and handling procedure from the learning days. Results were quantified with AnyMaze software and analyzed using GraphPad Prism.

### Western Blot

Prefrontal cortex, hippocampus, and cerebellum tissue samples were collected from both UT and COM rat brains. These samples were lysed using RIPA buffer with a protease inhibitor cocktail (ThermoFisher Cat# 89900 and 78440), and protein concentration was determined using Bradford assay. For western blots, equal amounts (30µg, unless stated otherwise) of protein samples from each group were electrophoresed on 10% SDS-PAGE gel under reducing conditions by beta-mercaptoethanol followed by transfer to PVDF (pore size 0.2µm) membranes. Blots were blocked with 5% nonfat dry milk in tris buffered saline with 0.075% tween 20 and probed with antibodies recognizing Aβ42. Monoclonal antibodiy recognizing the N-terminal (Abcam, cat#ab201060) of Aβ42 peptide was used along with Antibodies recognizing AQP4 (DIF8E, cell signaling), LYVE1 (rabbit, ThermoFisher), and GAD67 (mouse IgG2a, Millipore) were also used as described in the results section. The membranes were washed with 1X pbs and probed with secondary antibody goat anti-rabbit/chicken/mouse IgG (GeneTex). Chemiluminescence (Pierce, Rockford, IL) signals were detected by Flurochem M (Protein Simple, San Jose, CA) scanner. Analysis was performed in Graph pad prism.

### Proteomics

Hippocampus samples obtained from six UT and five COM rats were homogenized in RIPA buffer containing protease and phosphatase inhibitor (ThermoFisher, Cat# 78440). Protein samples were precipitated using methanol and then resolubilized in S-Trap loading buffer. Samples containing 100μg protein were all brought to equal volume. Cysteine disulfide bonds were reduced using dithiothreitol (DTT), free sulfhydryl groups alkylated with iodoacetamide (IAA), and unreacted IAA quenched with an excess of DTT. Protein was again precipitated using methanol and precipitate loaded onto S-Traps. Samples were digested using PierceTM Trypsin Protease, MS grade (ThermoFisher Scientific). Peptides were recovered from S-Traps by centrifugation. The samples were analyzed in duplicate (2 × 5 μg) using LC-MS/MS. LC-MS/MS was performed utilizing a Lumos Fusion Orbitrap (ThermoFisher Scientific). Data was analyzed using Proteome Discoverer 2.5 (ThermoFisher Scientific) and results exported to an Excel (Microsoft) spreadsheet. Untargeted and relative differences in protein abundances were identified with a false discovery rate (q-values) less than 0.01.

### Transcriptomics

RNA was extracted from the hippocampus, using the PureLink RNA Mini Kit. Extracted RNA samples were sequenced on the Illumina NovaSeq SP v1.5 300 cycle system at Virginia Tech sequencing center. Sequencing data (fastq files) were aligned to the Rattus norvegicus reference genome (Rnor_6.0) using the STAR v2.7.3a software package. STAR was run with runThreadN=8, outSAMtype=BAM SortedByCoordinate, quantMode=GeneCounts, outSAMstrandField=intronMotif, and outFilterIntronMotifs=RemoveNoncanonicalreads. Filtered and normalized gene expression levels were calculated from the aligned reads using HTSeq v.2.0.5. Differentially expressed genes were identified by linear modeling and Bayesian statistics using limma v3.58.1 for R v.4.3.2.

## Results

### Quantification of pressure applied during COM treatment

COM was performed with the nanosensor glove on Yg-rat as shown in Fig 1A-C. Pressure applied on the rat skull during COM treatment and duration of treatment were recorded as shown in Fig 1D and E. COM duration has a cumulative average of 3.12 ±0.0849 minutes across 7 treatment days of data collected. The force applied on the skull was quantified from day 4 COM treatment recordings and the average force 1.95 ±0.162N, was calculated from all Yg-COM group animals. Clinical observations made by the COM practitioner are provided in **S1 Table**. Observations made on Yg animals indicate that the six COM treated rats on day 1 of the experiment, prior to any treatment, had an absent, restricted, or compressed CRI indicating amplitude. By day 7 and upon completion of the final treatment, their individual CRIs were good or fully symmetric, indicating improved bone and dural membrane mobility and presumably improved lymphatic circulation. Similar results were found in Tg treated rats as well. The five Tg-COM group (TgF344 rats) on day 1 had pretreatment CRIs that were classified as restricted or absent. By day 7, the CRIs were reported to be good or full, as shown in the **S2 Table**.

**Fig 1.**
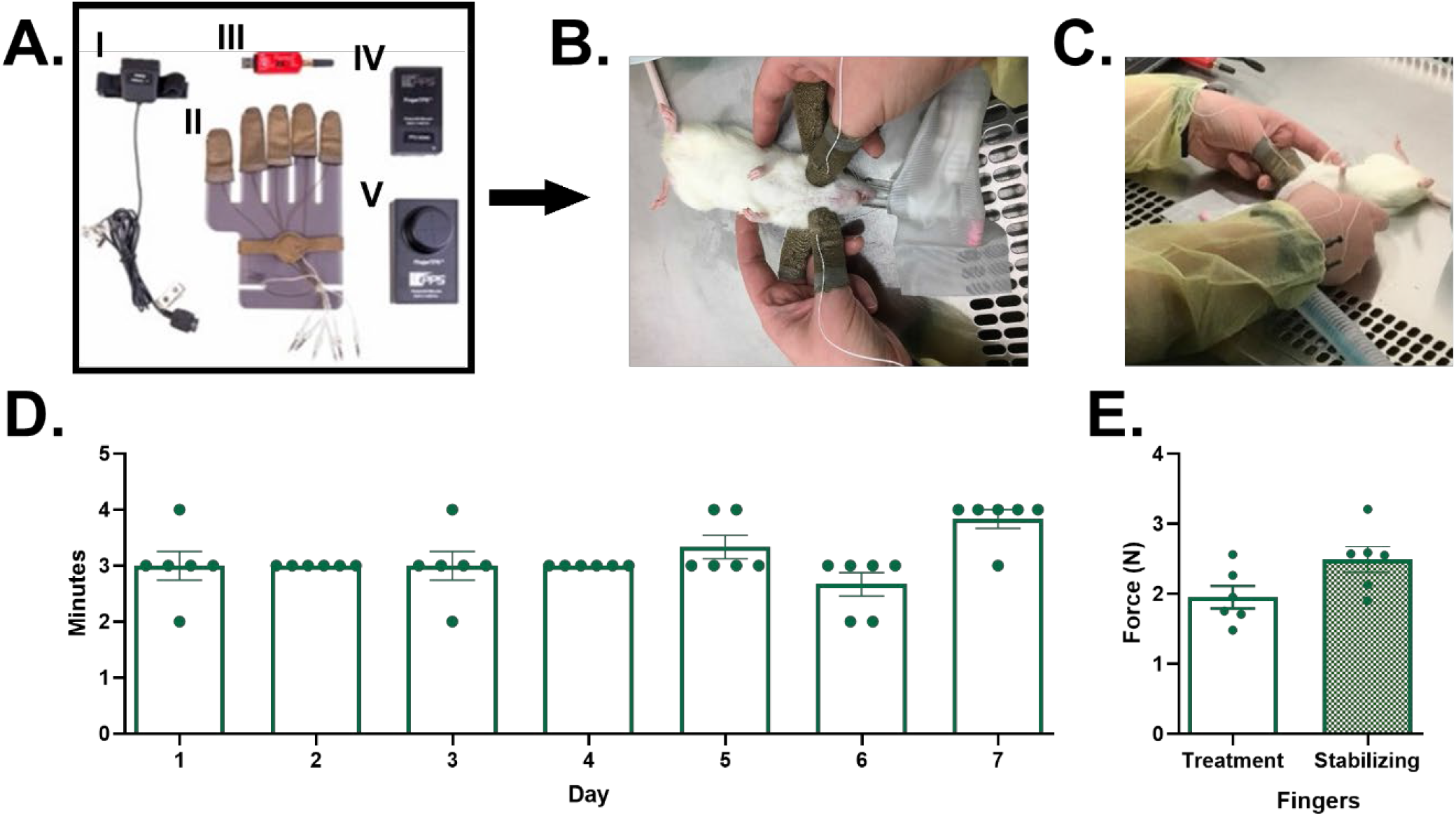
Pictures of components and use of FingerTPS pressure sensing device and associated preliminary data. **(A)**:1. Wrist mount and cable to sensor interface, 2. Custom made fingertip sensors, 3. Bluetooth dongle, 4. FingerTPS electronics, & 5. reference sensor. **(B)** Practitioner has worn sensors on stabilizing (pointer fingers) and treatment (middle) finger while treating a young adult rat to test device functionality and obtain preliminary data. **(C)** The picture shows the Bluetooth device connected to a computer to record forces generated by the fingers during COM procedure. Histograms represent the duration of COM treatment from preliminary phase 4 data set **(D)** for each animal, and average pressure applied by the treatment and stabilizing fingers on day 6, when acquisition was most accurate **(E)**. N=6 Yg-COM.

### COM Alters Spatial Learning and Memory Related Phenotypes in Rat Model of AD

Both Yg and Tg rats completed the MWM assay over the course of 8 days. For Yg, the learning days data showed significant reduction in escape latency when placed in the NW zone (Fig 2A). When comparing escape latency across the days and within groups, there was statistical significance of p=0.016 for the Yg-UT rats. Specifically comparing days 1 and 2 (45.7±6.63s vs 16.2±4.43s) and 1 and 4 (45.7±6.63s vs 5.50±2.35s) this highlights a decrease in escape latency. The Yg-COM rats’ data also show the similar trend through significant differences when comparing days 1 and 3 (60.0±0.00s vs 5.83±2.71s, p=<0.001) and 1 and 4 (60.0±0.00s vs 14.0±6.65s, p=0.024). When comparing the treatments, Yg-UT (17.2±3.98s) and Yg-COM (5.83±2.71s) within the days, a significance of p=0.044 was found on day 3. Passing of normality for the eight charts that were assessed with the Shapiro-Wilk Test. Comparisons were done using a repeated measures 2-way ANOVA or mixed effect model with Geisser-Greenhouse correction and a Tukey multiple comparison test. Escape latency for testing days 1-7 were analyzed using a 2way-ANOVA and uncorrected Fisher’s LSD (Fig 2B). An unpaired parametric t-test was used for day 8. Transformations were used when data sets did not pass the Shapiro Wilk normality test. Significance was only found for platform visits on day 8 with the Yg-UT rats having significantly more (6.17±0.910 vs 3.50±0.671, p=0.040). Fig 2C illustrates the performance of the rats on day 8 of the assay. Most of the parameters reported showed no significant differences except for time moving towards (Yg-COM 13.3±0.457s vs Yg-UT 15.4±0.353s, p=0.004) and away from the platform zone (Yg-COM 9.05±0.694s vs Yg-UT 11.5±0.590s, p=0.040). Overall, for Fig 2C, Yg-UT rats mostly have higher values compared to the Yg-COM rats. This set of analysis was also performed with an unpaired parametric t-test. Welch’s correction was used when the data set for the unpaired parametric t-test did not pass the F-test. Transformations and nonparametric tests were performed on data sets that did not pass the Shapiro Wilkes normality test.

**Fig 2.**
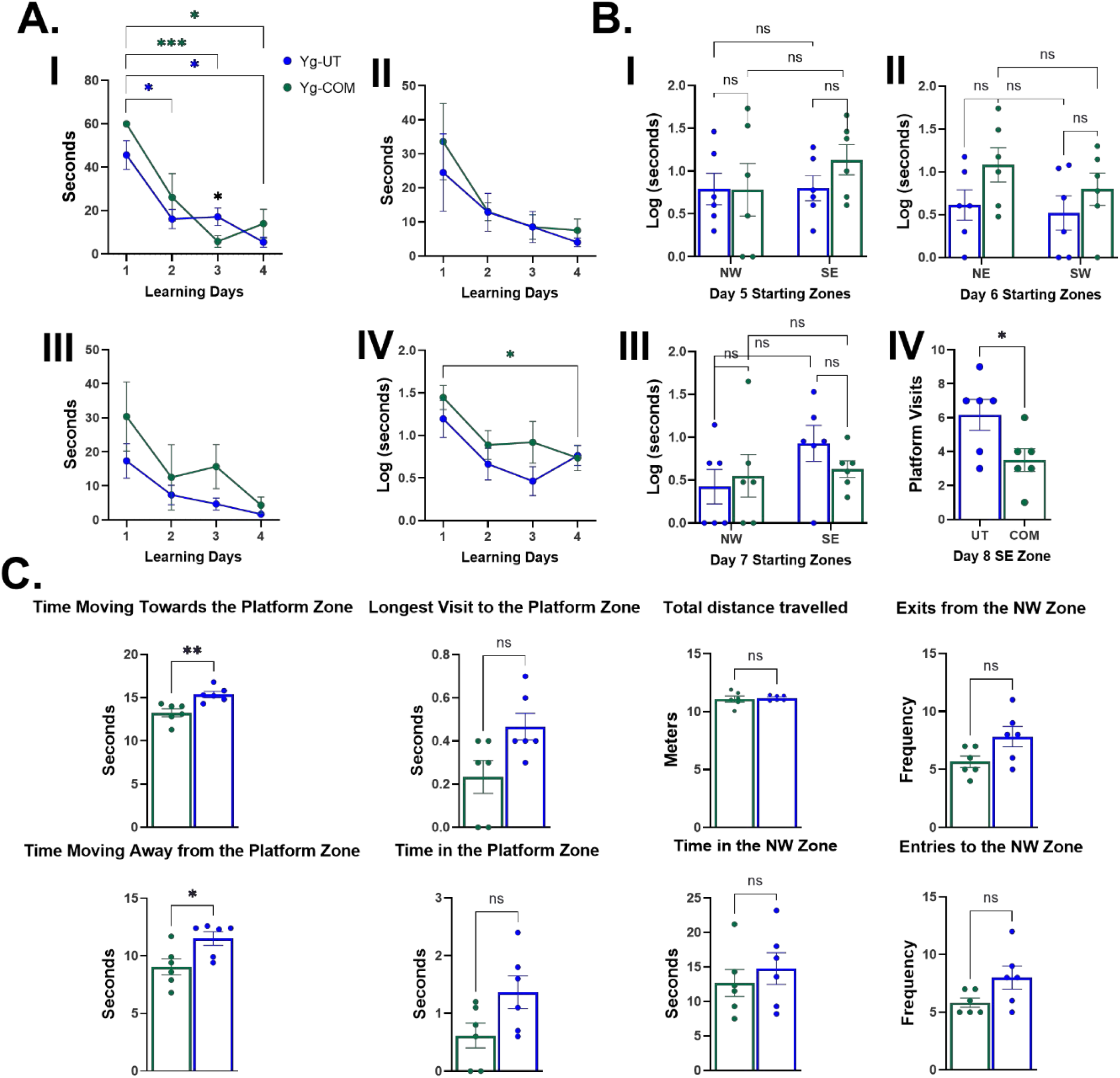
Wild type 3-month-old (Yg) Morris water maze manual and anymaze recording results. **(A)** Manual Recording Escape Latency from Northwest (I), Northeast (II), Southwest (III), and Southeast (IV) mixed effect analysis comparisons for learning days (1-4). **(B)** Manual Recordings for serial phase shift repeated learning multiple unpaired t-test comparisons for each testing day(I, II,III), along with platform-less day 8 unpaired t-test comparison(IV). Histograms I, II, and III were logarithmically transformed and the raw data is in **S3 Raw Fig. (C)** Day 8 parameters tracked with Anymaze for treatment unpaired t-test comparison of associated learning and memory behaviors. (N=12, 6 male and 6 female, 6 per group, APA p-value style: 0.12(ns), 0.033(*), 0.002(**),0.001(***), SEM reported.

In the Tg cohort learning trials, a significant difference was observed within the Tg-UT rat escape latency for days 1 and 4 when placed in the NE (38.7±8.94s vs 9.83±3.59s, p=0.032). Significance was also found in SE zone days 1 and 3 (47.8±8.25s vs 10.2±3.58s, p=0.024) and days 1 and 4(47.8±8.25s vs 17.5±5.86s, p=0.035) for Tg-UT in Fig 3A. No significance was found for the COM treated group across testing days and when starting in different quadrants. Comparisons were analyzed with a repeated measures 2-way ANOVA. Serial testing was also performed with rats being placed in a different quadrant for each of the testing days and no significance was found on days 5-7 for escape latency (Fig 3B). Unpaired parametric t-tests were used for analysis. Of the parameters studied using Anymaze, the histograms presented in Fig 3c were of interest and significant. Day 5 has the highest number of data sets (7 parameters) that illustrate significant difference between the two groups. Time moving towards the NW zone had a p-value of p<0.001 (Tg-COM 6.32±0.784s vs Tg-UT 17.4±1.70s), while the other 6 parameters had p-values less than 0.05. Day 6 has one being signed initial heading error to the platform zone (Tg-COM 35.2±23.8° vs Tg-UT -39.3±4.52°, p=0.034), while day 8 also has two. Latency to last entry to the platform zone has a p-value of 0.026 (Tg-COM 37.8±5.63s vs Tg-UT 12.2±5.67s) and path efficiency has p-value of 0.036 (Tg-COM 0.024±0.005 vs Tg-UT 0.068±0.015). Unpaired parametric and nonparametric t-tests were used. Unpaired parametric t-tests, that did not pass the F-test were analyzed with Welch’s correction. All figure 3 histograms were checked for normality with Shapiro Wilke’s test.

**Fig 3.**
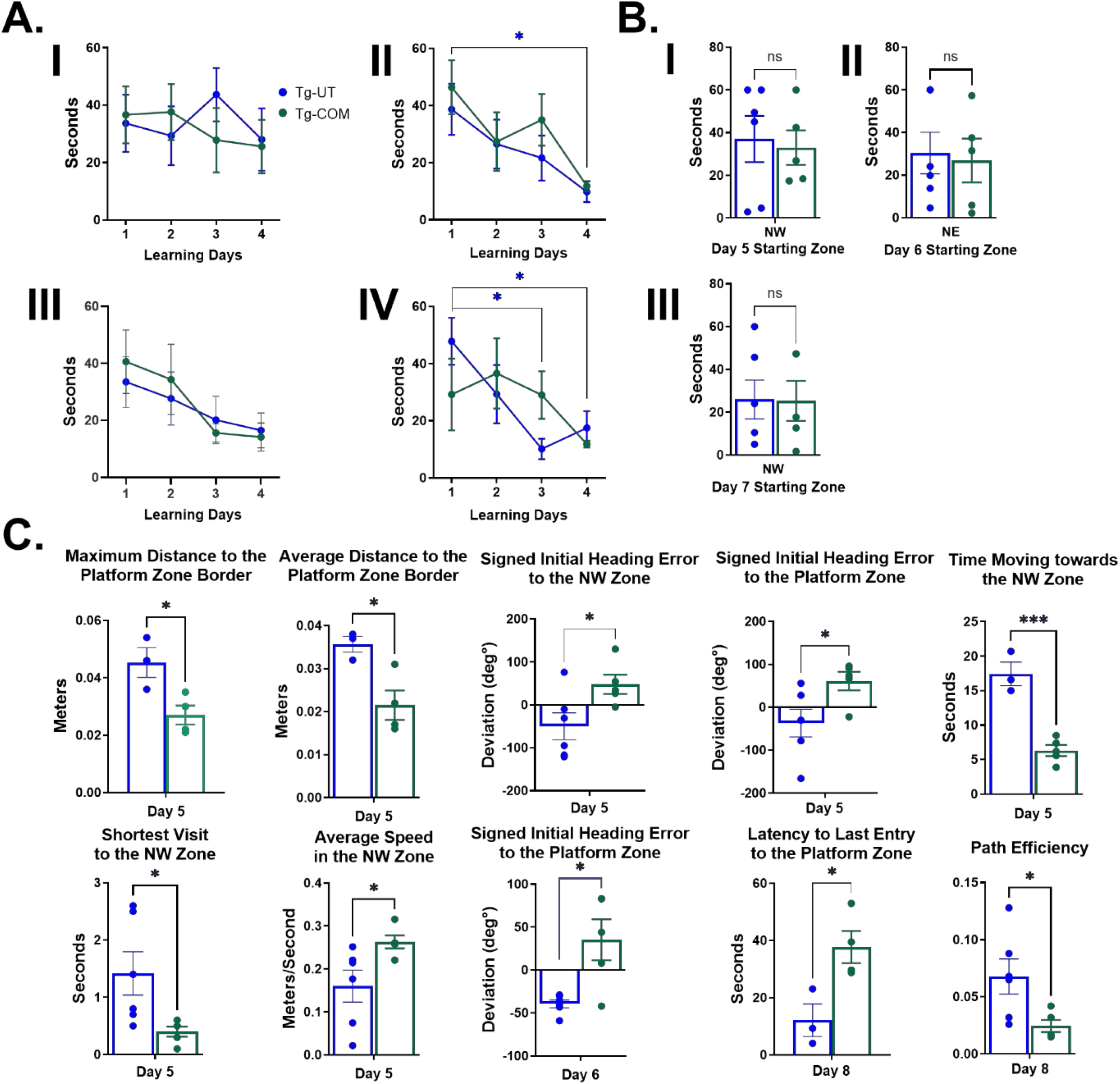
TG-F344 18-month-old (Tg) Morris water maze anymaze recording results. **(A)** Manual Recording Escape Latency from Northwest (I), Northeast (II), Southwest (III), and Southeast (IV) 2-way-ANOVA comparisons for learning days (1-4). **(B)** Manual Recordings for serial phase shift repeated learning unpaired t-test comparisons for each testing day(I, II,III). **(C)** Anymaze parameters associated with learning and memory that had significant differences between the treatments for each testing day. (N=11, 6 Tg-COM and 5 Tg-UT, APA p-value style: 0.12(ns), 0.033(*), 0.002(**),0.001(***), SEM reported.

### COM altered the expression of proteins associated with CNS fluid circulation in rat model of AD

Based on the previous reports (9,10,20), we choose to study the quantitative expression of proteins involved in the CNS fluid circulation and cognitive function using western blots; these proteins including Aβ42, AQP4, LYVE-1, and GAD67. The proteins of interest are depicted by their associated histogram for average blot expression. For Yg rats there were no significant differences between the two groups for each protein and brain region listed (Fig 4). For Tg-rats there was no significant differences between the groups for Aβ42 (Fig 5). However, GAD67 expression for Tg cerebellum was significantly increased for the Tg-COM groups (Tg-COM 1.24±0.118 vs Tg-UT 0.923±0.0838, p=0.49). Significance was also seen with LYVE-1 expression for P5 prefrontal cortex and hippocampus. The Tg-COM group had significantly higher expression of LYVE-1 in the prefrontal cortex (0.442±0.0234 vs 0.319±0.0159, p=0.002) and for the hippocampus the Tg-COM rats had significantly lower expression (0.208±0.0255 vs 0.281±0.0153, p=0.30). For both data sets, normality was checked with the Shapiro Wilk’s test. Comparisons were assessed with an unpaired parametric t-test; Welch’s correction was used when data sets did not pass the F-test.

**Fig 4.**
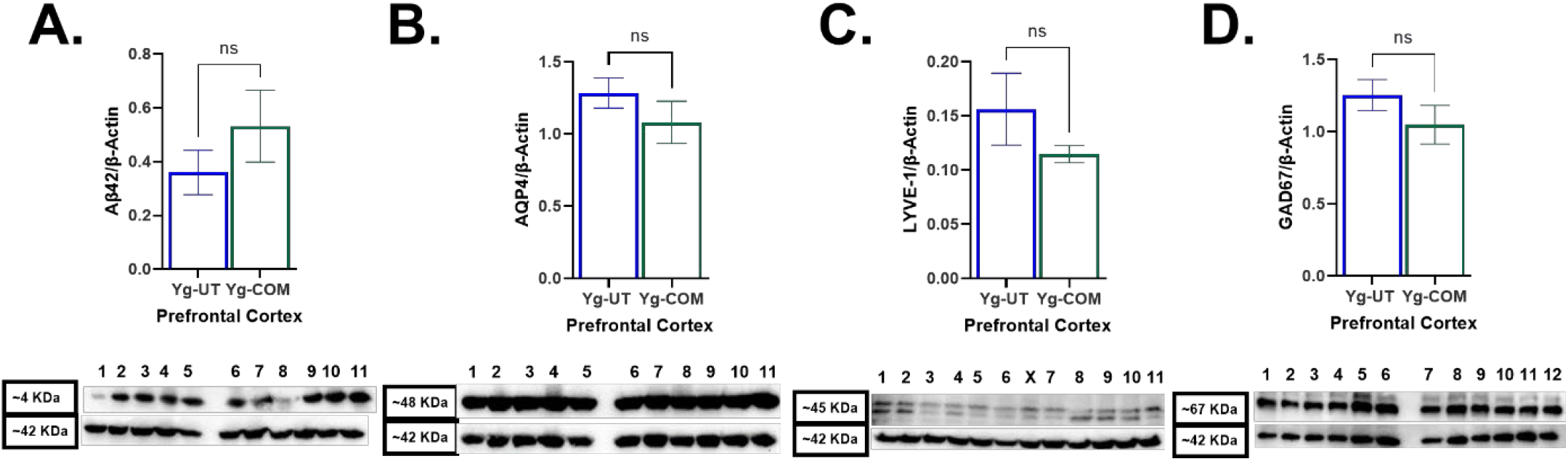
Western blot results from prefrontal cortex, hippocampus, and cerebellum for wild type 3-month-old (Yg). Representative western blot images, histograms of relative signal density, and unpaired t-test analysis for the expression of **(A)** Aβ42, **(B)** AQP4, **(C)** LYVE-1, and **(D)** GAD67. Numbers on the blot indicate UT (1-5) and COM (6-11) for A-C. For D, lanes 1-6 are UT and 7-12 are COM. Whole blot images are provided in **S4 Raw Images**. (APA p-value style: 0.12(ns), 0.033(*), 0.002(**), 0.001(***), SEM reported.

**Fig 5.**
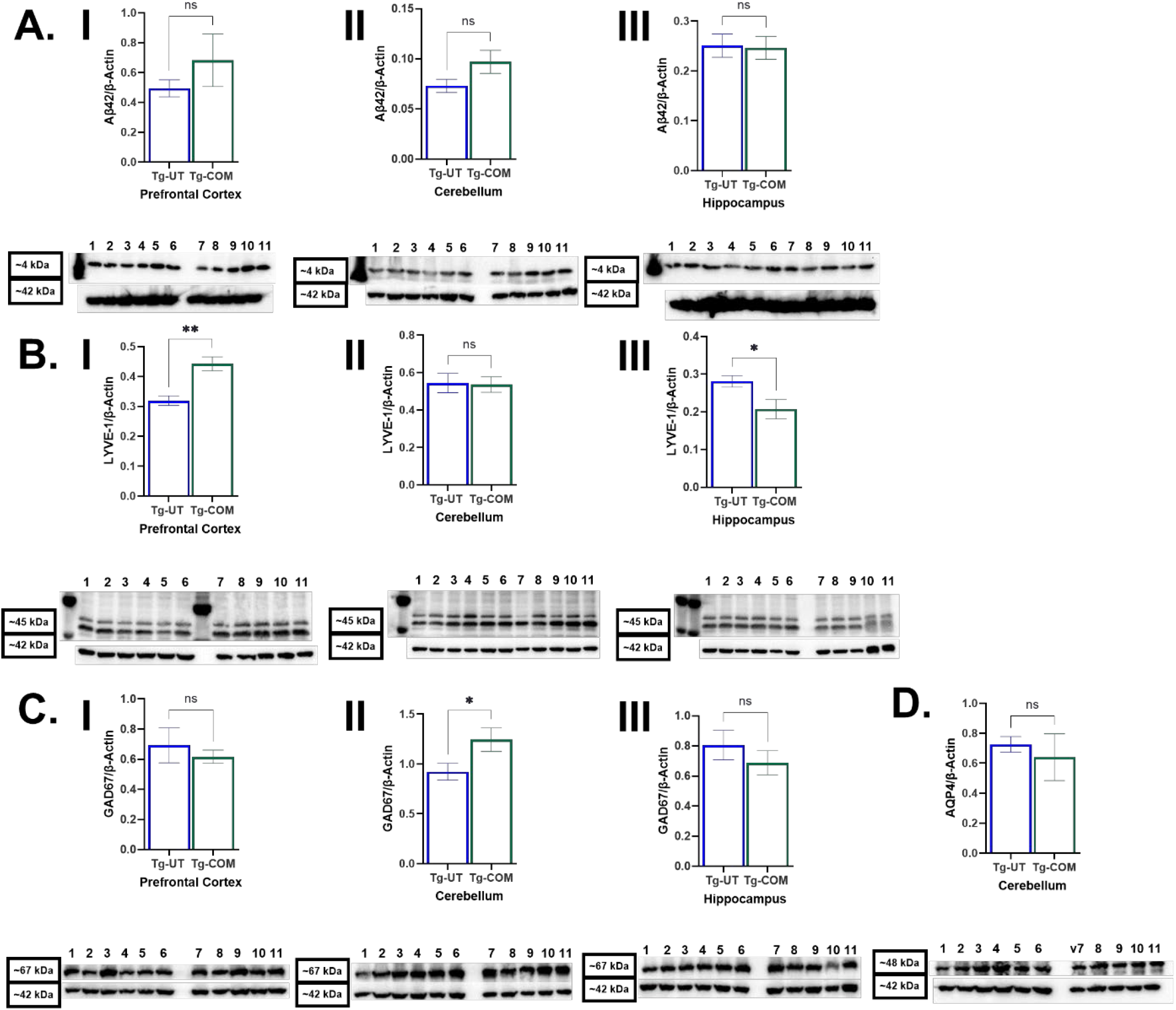
Western Blot Results from prefrontal cortex, hippocampus, and cerebellum for TG-F344 18-month-old (Tg). Representative western blot images, histograms of relative signal density and unpaired t-test analysis for the expression of **(A)** Aβ42, **(C)** LYVE-1, **(D)** GAD67, and **(B)** AQP4. Numbers on the blot indicate UT (1-6) and COM (7-11) samples. Whole blot images are provided in **S5 Raw Images**. (APA p-value style: 0.12(ns), 0.033(*), 0.002(**), 0.001(***), SEM reported.

To identify the COM induced changes in protein expression, a proteomic assay was performed using hippocampus, a region primarily involved in learning and memory. False Discovery Rate confidence was set to 0.01 which identified 86 differentially expressed proteins. Fold change values show 33 proteins upregulated and 53 down regulated (Fig 6). To perform a more rigorous analysis mascot scores were also set to filter out all values less than 100 or blank. This left 45 different proteins that COM significantly altered the expression. Of the 45, 34 proteins are associated with neurological disorders and diseases including Serine-threonine kinase P21-activated kinase 3 (PAK3). A table of these proteins and the generally associated disorder can be found in **S6 Table**. A transcriptome analysis was also performed but that did not identify any significant differentially expressed genes. The transcriptome figure can also be found in **S7 Fig**.

**Fig 6.**
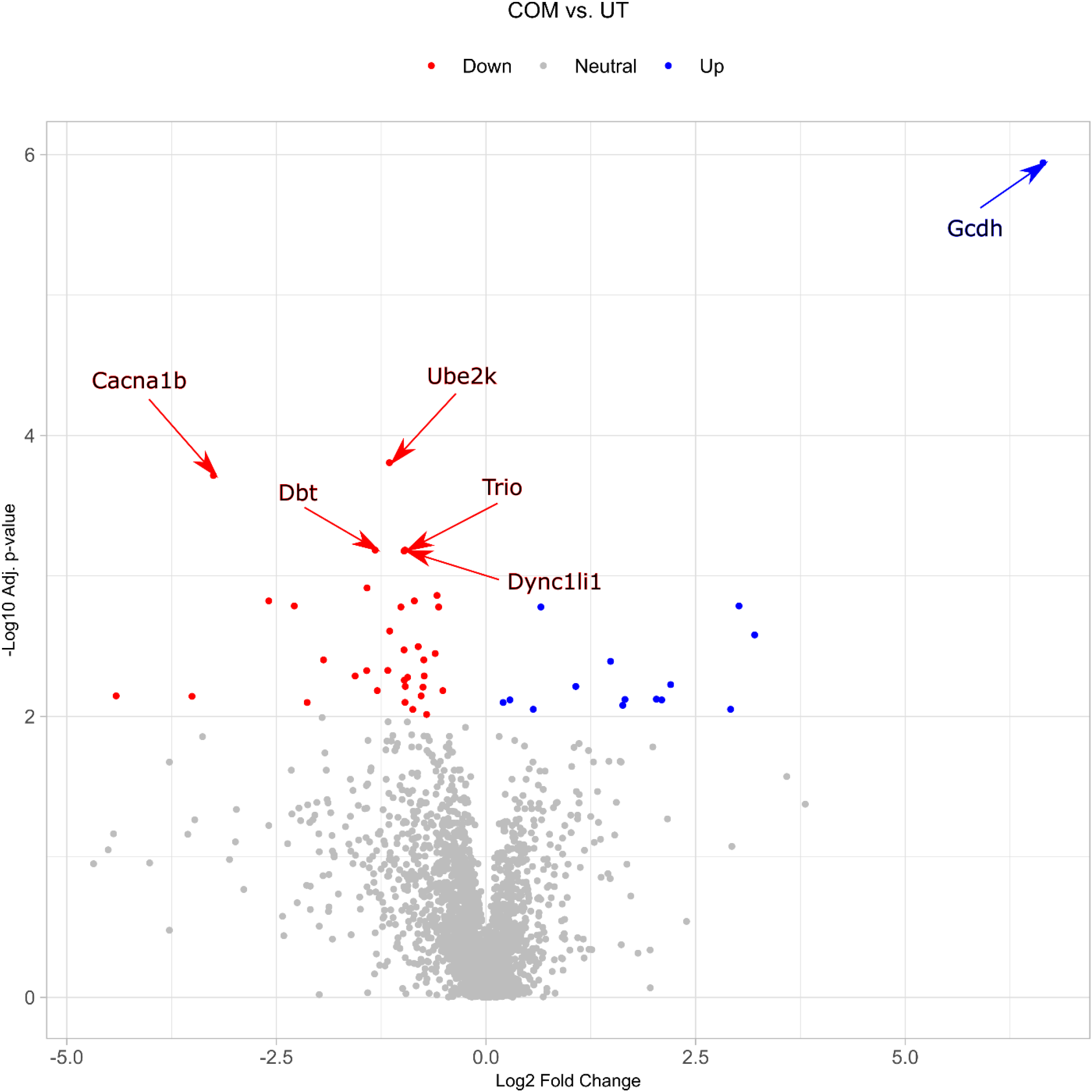
Proteomic analysis of TG-F344 18-month-old (Tg) rat brain hippocampal tissue. Volcano plot shows differential expression proteins in COM treated rats compared to the UT group. The analysis identified that COM significantly up **(blue)** and down **(red)** regulated the expression of 45 proteins including 34 Alzheimer disease related proteins. False Discovery rate (q-values) set to less than 0.01. False Discovery rate (q-values) set to less than 0.01.

## Discussion

In this study, for the first time we have established a technique to quantify the force applied during COM treatment on an animal model of AD. This quantifiable COM treatment technique will be useful for future studies on AD and other relevant animal models.

Results from the Yg cohort indicated that COM does not pose a significant effect on phenotypes associated with AD. Across all days of testing both Yg-COM and Yg-UT rats performed relatively the same with both showing a trending decrease in escape latency. However, on day 8, Yg-UT rats visit the platform zone more frequently than the Yg-COM treated rats (Fig 2B). This could be an indicator of exploration willingness, poor performance, or nonperformance of the Yg-COM group rats. Most of the day 8 parameters show no difference and the two that do are, time moving towards and away from the platform zone (with average values significantly lower for the Yg-COM rats. This might suggest that once the escape route was removed, the Yg-COM rats were more likely to explore the escape possibilities compared to their counterparts. Alternatively, development of learned helplessness, a hippocampus independent learning, could also be a valid consideration(21). Since the rats participated in MWM over the course of multiple days, repetitiveness could have given them security in the platform always being present. Once removed, escape was no longer an option so, the rats could opted to “give up” and float, explore other areas, or continue to check the same area over and over for the missing platform(22).

Therefore, an interpretation that COM treatment improved exploration in Yg rats could be given. These results also indicated that COM has no negative impact on the spatial learning and memory-related parameters studied on Yg rats.

In the Tg-cohort, however, multiple parameters were different between the Tg-UT and Tg-COM groups unlike the Yg cohort. Despite the indifference in escape latency, Tg-COM rats swam faster and were located closer to the platform compared to the Tg-UT rats (Fig 3). Even though the Tg-UT rats have a more efficient path by day 8 to reach the location where the platform used to be, this does not indicate if the path taken was hippocampus dependent or independent. However, the heading error, which could be interpreted as an effective approach to reach the platform, is statistically different between the groups. Tg-COM rats had a positive deviation meaning the target zone is to the animal’s right, while Tg-UT rats had a negative deviation, indicating the target zone is to their left. This might contribute to the Tg-COM group’s lower path efficiency. In future works, we could characterize identify specific hippocampus dependent paths which would be helpful to distinguish spatial learning behaviors.

Results from the immunoassays validate findings from behavioral assays since there lacks significance in the expression of proteins associated with CNS fluid circulation (Fig 4 and 5). There was no difference in the expression of Aβ42, AQP4, LYVE-1, and GAD67 between the Yg-COM and Yg-UT. This further suggests that COM poses no significant adverse risk or effects on a healthy young rat brain. Further, in the P5 cohort, indifferent expression of Aβ levels suggest that the number of COM treatments in the Tg rats was not enough to see a difference. However, previous studies have reported a reduction in Aβ levels in 18-month old wild type rats(9). This could be due to blockade from the amount of Aβ accumulation or plaque formation specific to the genetically modified Tg rats. However, the proteomic analysis identified 45 proteins differentially expressed in COM treated hippocampus. Of these proteins, 34 are associated with neurological disorders such as AD dementia, seizure disorders, movement disorders, and psychiatric disorders. The most significantly upregulated protein was PAK3 which is responsible for regulation of neuronal synaptic morphology, functionality, and outgrowth(23). In the AD brain, PAK3 is typically reduced or dysregulated which can lead to developmental cognitive deficits(23,24). Since PAK3 is upregulated in the Tg-COM rats, this suggests that COM could have neuroprotective effects through modulation of PAK3. However, this needs to be validated by more quantitative immune assays.

In contrast, the transcriptomic assay performed in this study identified no significant (FDR < 0.01) differentially expressed genes in COM treated rats. We recognize that the correlation between mRNA and protein expression is not always consistent as previously reported(25,26). Generally, proteins have a longer half-life than mRNAs. Thus, proteins that are synthesized after every COM treatment might not have degraded before the next COM treatment. Therefore, there might not be enough mRNA trigger to produce more protein while already synthesized proteins are sufficient to carry out the cellular function that needed to be performed in response to the COM treatment. This is in agreement with the interpretation that the variability of the mRNA expression is indicative of the cell controlling mRNA expression at different points of the cell cycle to achieve the desired protein levels(27).

However, the transcriptome results were also inconsistent with our previous work on wild type (WT) rats(20), where 688 genes were differentially expressed in COM treated rats compared to WT rats, with FDR < 0.01. There are several possible reasons for the differing results. One possibility is that different rat strains were used in the two studies. In the previous study, we used a WT rat model (aged rats) as opposed to the Tg rats used here. Multiple genetic mutations to induce the pathophysiology of Alzheimer’s disease might have altered the normal mRNA and protein synthesis regulatory mechanisms. It is also possible that COM treatment could not affect the mRNA expression in the Tg rats, that have pathogenesis evolving from the nucleic acid level used in this study, as much as the naturally aged 18 months old rats used in the previous study. Nonetheless, it was sufficient to affect protein expression in both cases. Another possibility is that hippocampal tissue was used in this study instead of the prefrontal cortex used previously. The effect of COM treatment may be less pronounced in the hippocampus compared to the prefrontal cortex, but sufficient to affect protein expression.

In summary, we have successfully performed a quantifiable COM on Yg and Tg rats and compared its effect on AD-relevant phenotypes for the first time. COM produced more behavioral and biochemical changes in aged Tg rats than in Yg rats. Remarkably, COM influenced spatial learning behaviors and cognition in a Tg rat model by altering proteins involved in CNS fluid circulation and cognitive function.

### Limitations

The findings in this study should be considered in light of potential limitations. The cohort sizes are large enough for us to achieve power for each phase, but they were also too small to remove outliers and investigate sex differences. Currently, with the extramural funding, we are investigating the sex differences, as Alzheimer’s disease is known to predominately effect women.(28)

## Ethics Declaration

All experimental procedures were conducted in accordance with the Institutional Animal Care and Use Committee (IACUC) of Virginia Tech (Protocol ID# 15-099 and ID#19-045).

## Acknowledgements

Work was funded by grant# 1915733 American Osteopathic Association (AOA) and grant# 1R15 AT010789-01A1 National Institute of Health (NIH) to Blaise Costa. Authors acknowledge Costa lab research assistants and animal house staff for their help with the MWM assay.

## Author Contribution

BMC performed behavioral experiments, biochemical assays, analyzed data, and wrote the manuscript. DH analyzed data and wrote the manuscript. HT performed COM treatments for both cohorts. PD analyzed MWM data. SB performed Western Blot assay and analysis. RH performed proteomics and analyzed data. RA analyzed transcriptome and proteome data and interpreted the results. PV trained, provided scientific advice to DH, and reviewed manuscript.

## Data Availability Statement

All data associated with the results presented in this manuscript are provided in the figures and supplemental files. Raw data can be obtained from the corresponding author with a reasonable request.

## Supplemental

**S1 Table. Wild type 3-month-old (Yg) cranial manipulation clinician notes**. Physician documentation of duration and cranial rhythmic impulse for cranial manipulation treatment days 1-7.

**S2 Table. TG-F344 18-month-old (Tg) cranial manipulation clinician notes**. Physician documentation of duration and cranial rhythmic impulse for cranial manipulation treatment days 1-7.

**S3 Raw Fig. Wild type 3-month-old (Yg) Morris water maze raw manual recordings**. Raw data prior to normalization transformation of serial phase shift repeated learning histograms for each testing day.

**S4 Raw Images. Wild type 3-month-old (Yg) whole western blot images**. Annotated images of uncropped western blots for (A) Aβ42, (B) AQP4, (C) LYVE-1, and (D) GAD67 expression.

**S5 Raw Images. TG-F344 18-month-old (Tg) whole western blot images**. Annotated images of uncropped western blots for (A) Aβ42, (B) LYVE-1, (C) GAD67, and (D) AQP4 expression.

**S6 Table. Alzheimer’s Disease and Neurological disorder proteomic list**. Significantly differentially expressed proteins associated with neurological disorders from Tg rat brain hippocampal tissue proteomic analysis.

**S7 Fig. Transcriptomic analysis of TG-F344 18-month-old (Tg) rat brain hippocampal tissue**. Volcano plot shows differential expression proteins in COM treated rats compared to the UT group.

