## Supplemental document 1 for "Effect of quantified cranial osteopathic manipulation on wild type and transgenic rat models of Alzheimer’s disease"

### Rat Trial #4 Clinical Observations January 2020

| 3mo Wild Rats . Overall very small – size made them hard to treat. I am not sure treatments were the most effective but they were not as rigid as older Rats | **3 male** | **6 male** | **4 male** | **12 female** | **13 female** | **14 female** |
|---|---|---|---|---|---|---|
| **1/3/2020**<br><br>**Treatment Duration** | CRI Absent - Compression<br>3:30 | Mild Compression<br>CRI restricted<br>3:19 | CRI restricted Rt<br><br>2:00 | CRI Absent<br><br>3:10 | CRI absent<br><br>3:10 | CRI Absent<br><br>4:00 |
| **Post Treatment** | CRI Increased | Inc CRI but still Comp | ICRI More Symmetric | Inc CRI still Mod Comp | Inc CRI Poor Decreased Rt | CRI Poor Decreased Lt |
| **1/4/2020**<br><br>**Treatment Duration** | CRI absent - Compression<br>3:00 | CRI Poor<br><br>3:?? Timing Error | CRI almost absent<br><br>?:?? Timing Error | CRI Absent Compress<br><br>3:38 | CRI poor Decreased Rt<br><br>3:22 | CRI Absent Compress<br><br>3:16 |
| **Post Treatment** | CRI Improved and more symmetric | CRI Fair | CRI Poor | CRI still Mod Compression | CRI Fair more symmetric | CRI Poor |
| **1/5/2020**<br><br>**Treatment Duration** | CRI absent - Compression<br>3:38 | CRI Poor<br><br>3:21 | CRI Poor<br><br>3:32 | CRI Absent Compress<br><br>3:08 | CRI poor Decreased Rt<br><br>2:50 | CRI Absent Compress<br><br>4:30 |
| **Post Treatment** | CRI increased but poor | CRI Good Symmetric | CRI Fair | CRI Poor | CRI Fair | CRI Improved but poor |
| **1/6/2020**<br><br>**Treatment Duration** | CRI absent – Compression<br>3:25 | CRI Poor<br><br>3:18 | CRI Poor<br><br>?:?? Timing Error | CRI Absent<br><br>3:02 | CRI poor<br><br>3:10 | CRI Poor Decreased Lt<br>3:?? Timing Error |
| **Post Treatment** | CRI Poor | CRI Fair | CRI Fair | Energy Ball release CRI Poor | CRI Good | CRI Fair Decreased Lt |
| **1/7/2020**<br>**Treatment Duration** | CRI Poor<br>3:00 | CRI Fair<br>3:35 | CRI Poor<br>4:10 | CRI Poor Decreased Lt<br>3:02 | CRI Poor Decreased Rt<br>3:45 | CRI Poor Decreased Lt<br>4:00 |
| **Post Treatment** | CRI Fair | Bouncing still point CRI Good | CRI fair | CRI Good Symmetric | CRI Fair | CRI Fair symmetric |
| **1/8/2020**<br>**Treatment Duration** | CRI Fair<br>3:24 | CRI Good<br>3:14 | CRI fair<br>2:30 | CRI Fair<br>2:45 | CRI Poor<br>3:02 | CRI fair Poor<br>3:23 |
| **Post Treatment** | CRI good | CRI Full | CRI Good | CRI Good | CRI good | CRI good |
| **1/9/2020**<br><br>**Treatment Duration** | CRI restricted<br><br>4:30 | CRI Fair slight Decreased Rt<br>4:25 | CRI Good<br><br>4:00 | CRI Fair<br><br>4:30 | CRI Fair<br><br>4:00 | CRI Poor<br><br>4:30 |
| **Post Treatment** | CRI Good | CRI Good | CRI Full Symmetric | CRI Good | CRI Good | CRI Good |

OCC = Occipital Condylar Compression Rt = Right Lt = Left
