## Supplemental document 2 for "Effect of quantified cranial osteopathic manipulation on wild type and transgenic rat models of Alzheimer’s disease"

### Rat Trial #5 Clinical Observations June 2020

| **Cage**<br>**Marking**<br>**Sex** | **3**<br>**3L**<br>**F** | **4**<br>**4R**<br>**F** | **4**<br>**4L**<br>**F** | **2**<br>**2RR**<br>**M** | **2**<br>**2L**<br>**M** |
|---|---|---|---|---|---|
| **Day 1**<br>**Treatment Duration**<br>**TPS sensor pressure** | CRI Restricted. Decreased Lt<br>3:10<br>120-130g | CRI absent. Compressed<br>3:35<br>110-130g | CRI absent. Compressed<br>3:42<br>120-220g | CRI absent. Compressed<br>3:14<br>180-220g | CRI absent. Compressed<br>3:12<br>110-120g |
| **Post Treatment** | CRI Increased - fair<br>Still Decreased more on Lt | CRI increased - fair | CRI increased – poor<br>Decreased more on Rt | CRI increased – poor<br>Decreased more on Lt | CRI increased – poor<br>Decreased more on Rt. Rt OCC |
| **Day 2**<br>**Treatment Duration**<br>**TPS sensor pressure** | CRI absent. Compressed<br>3:48<br>120-250g | CRI Poor Mild Compression. Rt OCC<br>3:20<br>130-150g | CRI absent. Compressed<br>4:10<br>90-150g | CRI absent. Compressed<br>3:00<br>130-140g | CRI Poor. Compression. Decreased more on Rt.<br>3:20<br>150-220g |
| **Post Treatment**<br>Had Blood Draw in AM | CRI increased – poor<br>Decreased more on Rt<br>More restricted than Day 1 | CRI increased – good<br>Decreased more on Rt | CRI increased – poor | CRI increased – fair<br>Decreased more on Lt | CRI increased – fair<br>Decreased more on Rt |
| **Day 3**<br>**Treatment Duration**<br>**TPS sensor pressure** | CRI poor. Decreased Rt.<br>3:35<br>150-220g | CRI poor. Decreased Rt<br>3:02<br>130-220g | CRI fair. Decreased Lt<br>2:45<br>130-150g | CRI absent. Compressed Rt OCC<br>2:52<br>120-140g | CRI Poor Mild Compression. Decreased Rt.<br>2:48<br>200-220g |
| **Post Treatment** | CRI increased – fair | Quivering still point.<br>CRI increased – fair | CRI increased – good | CRI increased – good | CRI increased – fair<br>Decreased more on Rt |
| **Day 4**<br>**Treatment Duration**<br>**TPS sensor pressure** | CRI fair. Decreased Lt<br>3:40<br>90-120g | CRI fair.<br>3:08<br>110-150g | CRI fair. Decreased Lt<br>3:10<br>150-180g | CRI poor. Rt OCC<br>3:30<br>TPS sensor malfunction | CRI poor<br>2:54<br>TPS sensor malfunction |
| **Post Treatment** | CRI increased – good<br>Decreased Lt | CRI increased - Full | Bouncing still point.<br>CRI increased – good | CRI increased – fair<br>Decreased Rt | CRI increased – fair<br>Mildly Decreased Rt |
| **Day 5**<br>**Treatment Duration**<br>**TPS sensor pressure** | CRI Good<br>3:00<br>110-130g | CRI Good<br>3:10<br>100-160g | CRI Good<br>2:35<br>100-160g | CRI fair. Decreased Rt<br>2:25<br>100-140g | CRI fair.<br>2:12<br>110-130g |
| **Post Treatment** | CRI increased – almost full<br>Mildly Decreased Rt | CRI increased - Full | CRI increased - Full | CRI increased – good<br>Decreased Rt. Rt OCC | CRI increased – almost Full |
| **Day 6**<br>**Treatment Duration**<br>**TPS sensor pressure** | CRI Good<br>2:48<br>120-160g | CRI fair. Decreased Lt<br>3:07<br>110-130g | CRI fair.<br>3:30<br>150-170g | CRI Poor. Decreased Rt<br>3:07<br>130-150g | CRI fair. Decreased Rt<br>2:58<br>120-140g |
| **Post Treatment** | CRI increased – almost full<br>Mildly Decreased Rt | CRI increased – good | CRI increased – good | CRI increased – Fair | CRI increased – good |
| **Day 7**<br>**Treatment Duration**<br>**TPS sensor pressure** | CRI Good. Decreased Lt.<br>2:35<br>120-130g | CRI Poor<br>3:20<br>130-200g | CRI fair<br>3:50<br>160-200g | CRI Good<br>3:20<br>130-170g | CRI fair. Decreased Rt<br>3:30<br>130-170g |
| **Post Treatment** | CRI increased - Full | CRI increased – good | CRI increased – good | CRI increased - Full | CRI increased – good |

OCC = Occipital Condylar Compression Rt = Right Lt = Left
