## Supplementary figures and images for "Effect of quantified cranial osteopathic manipulation on wild type and transgenic rat models of Alzheimer’s disease"

### Supplemental document 3

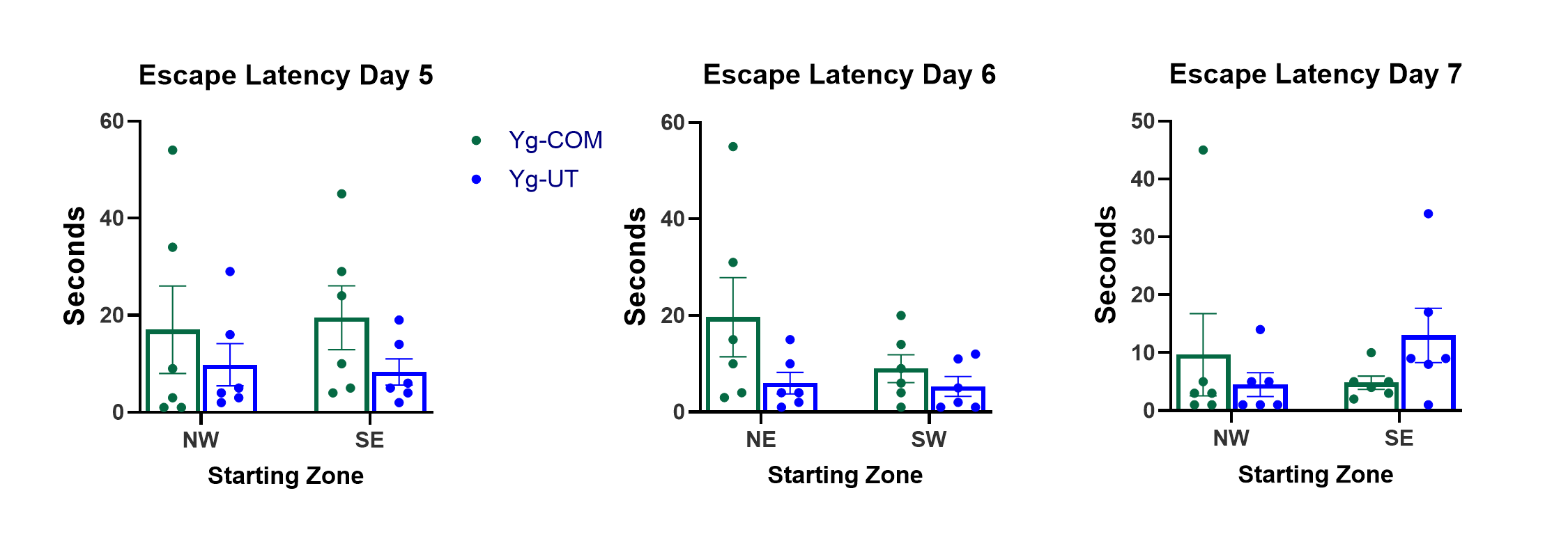

### Supplemental document 7

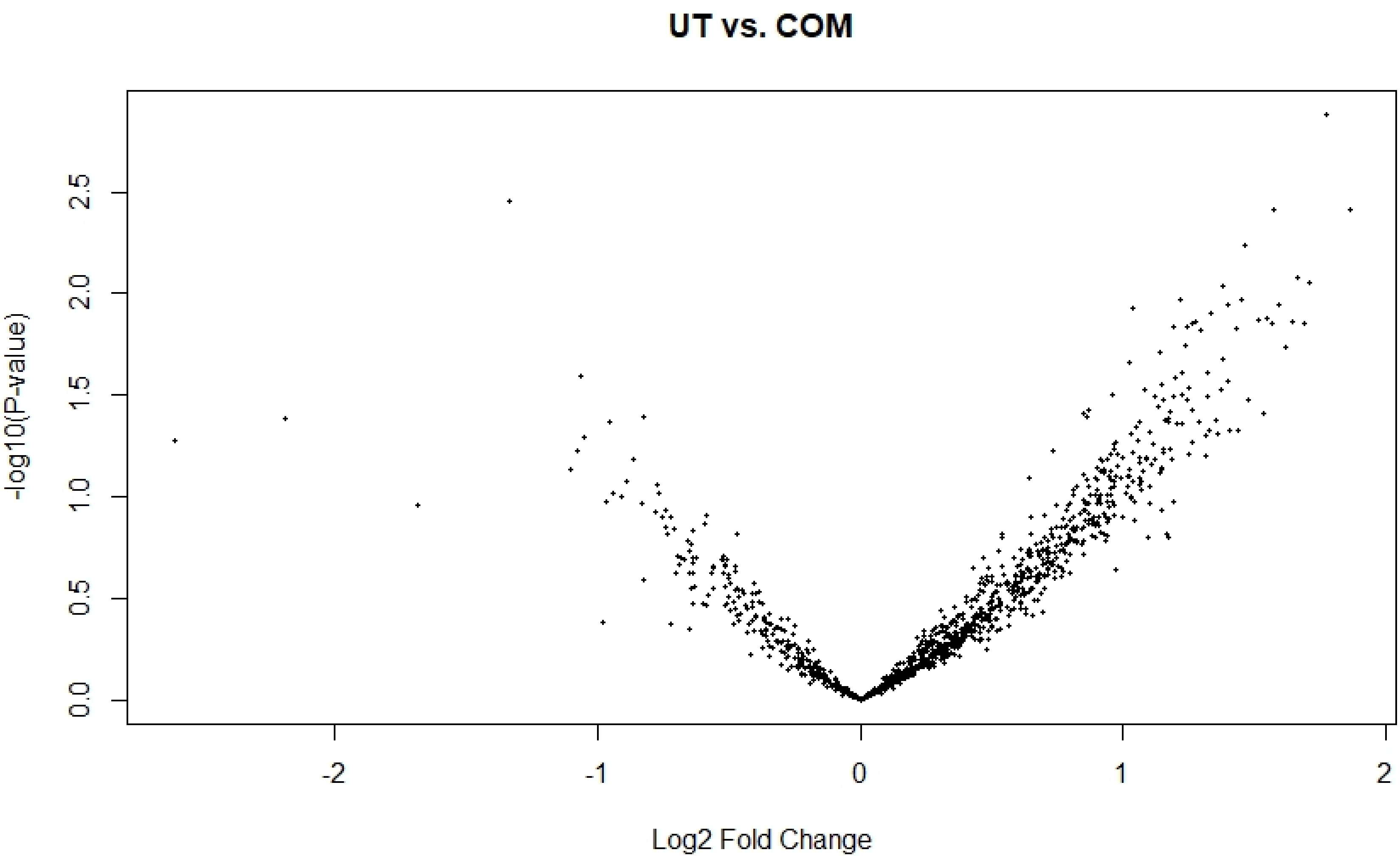
