## Supplemental Figure 4 for "Effect of quantified cranial osteopathic manipulation on wild type and transgenic rat models of Alzheimer’s disease"

**Supplemental 4 Raw Images. Wild type 3-month-old (Yg) whole western blot images.**
Annotated images of uncropped western blots for (A) Aβ42, (B) AQP4, (C) LYVE-1, and (D) GAD67 expression

Method to capture the following images: Chemiluminescence (Pierce, Rockford, IL) signals were detected by Flurochem M (Protein Simple, San Jose, CA) scanner per manufacturer guidelines.

**A) Antibody:** N-terminal, Abcam, cat#ab201060

*Prefrontal Cortex Aβ42*

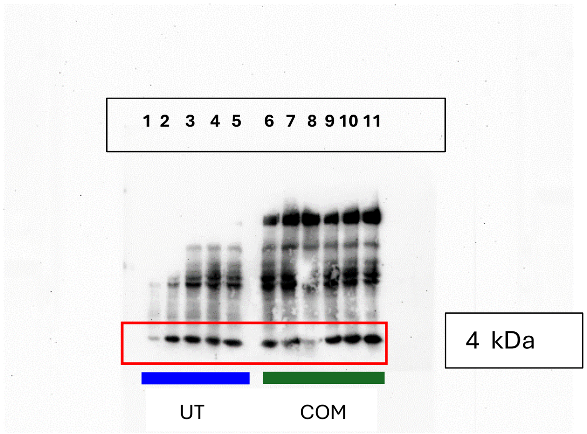


*β-Actin*

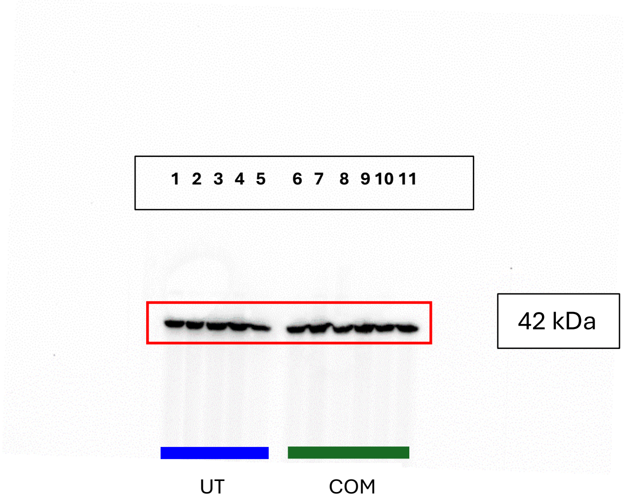

**B) Antibody:** DIF8E, Cell Signaling

*Prefrontal Cortex AQP4*

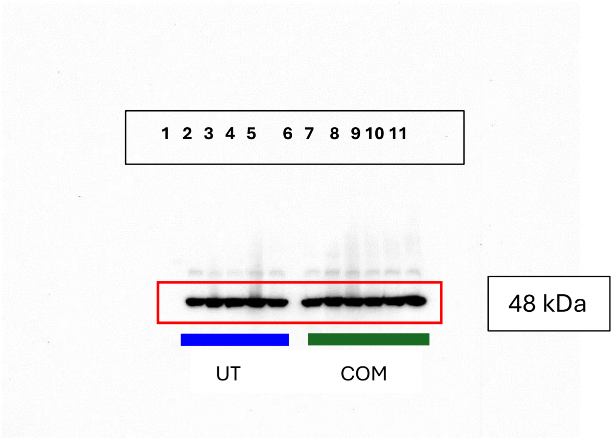


*β-Actin*

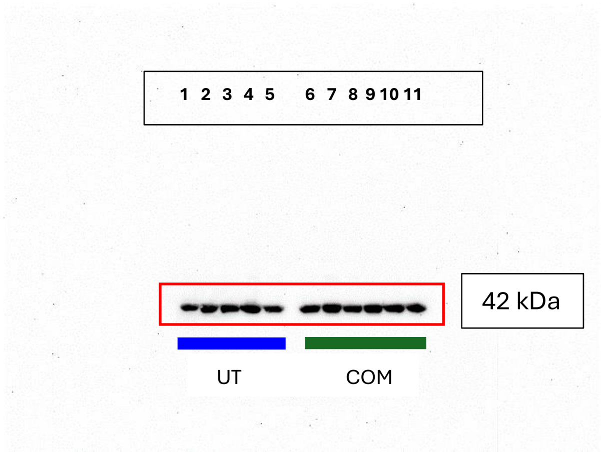

**C) Antibody:** rabbit, ThermoFisher

*Prefrontal Cortex LYVE-1*

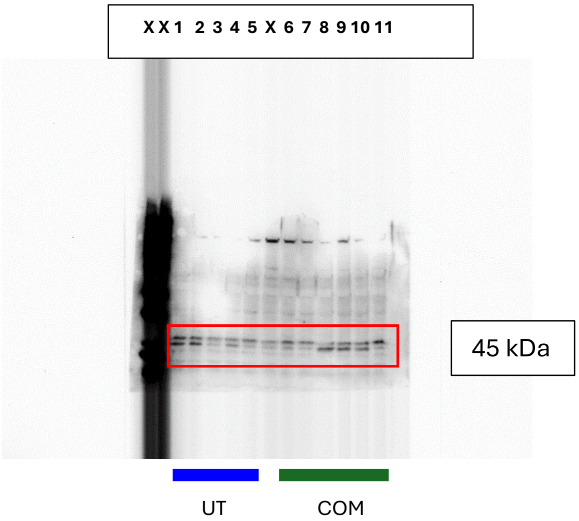


*β-Actin*

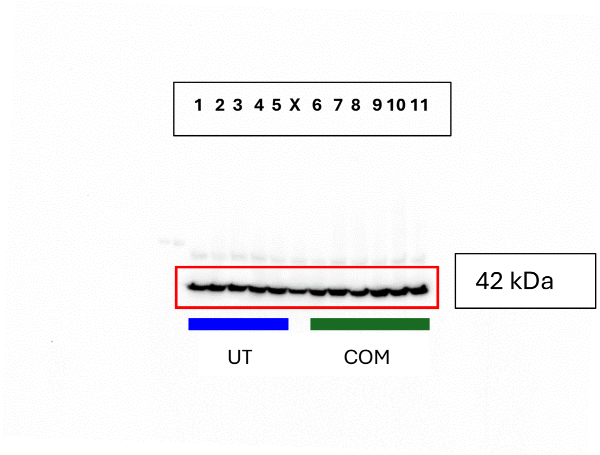

**D) Antibody:** mouse IgG2a, Millipore

*Prefrontal Cortex GAD-67*

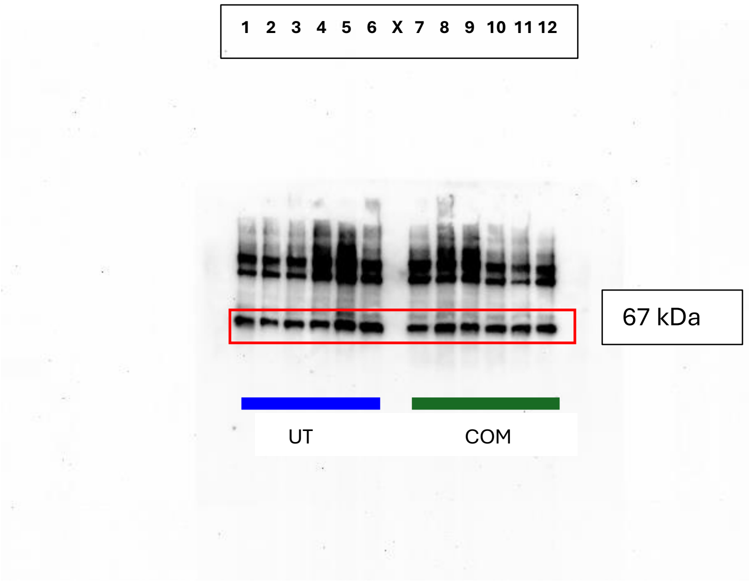


*β-Actin*

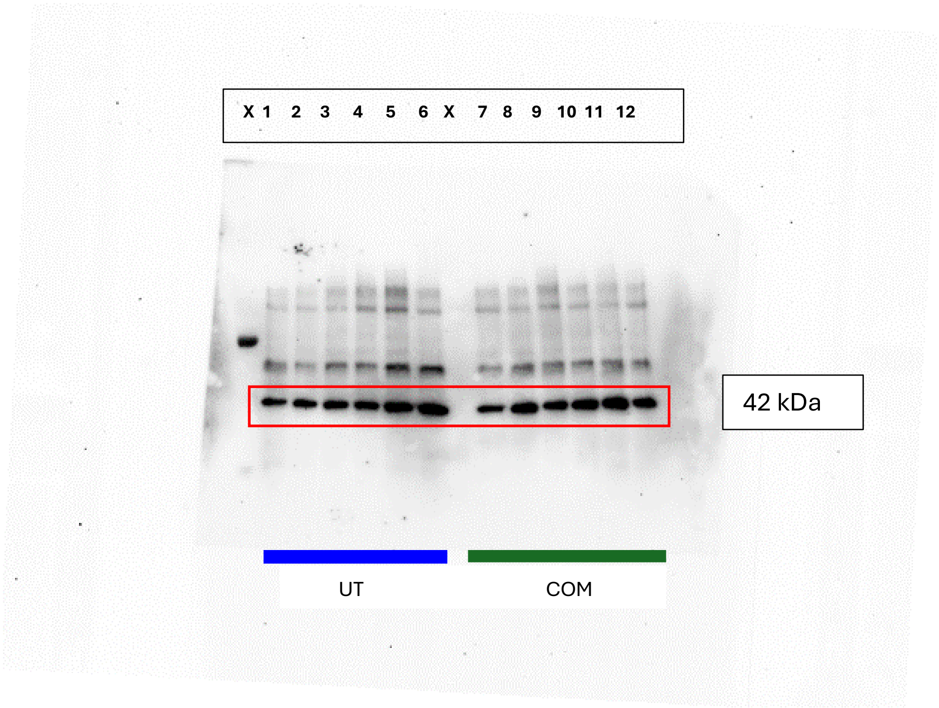
