## Supplemental document 5 for "Effect of quantified cranial osteopathic manipulation on wild type and transgenic rat models of Alzheimer’s disease"

**Supplemental 5 Raw Images. TG-F344 18-month-old (Tg) whole western blot images.** Annotated images of uncropped western blots for (A) Aβ42, (B) LYVE-1, (C) GAD67 and (D) AQP4 expression

Method to capture the following images: Chemiluminescence (Pierce, Rockford, IL) signals were detected by Flurochem M (Protein Simple, San Jose, CA) scanner per manufacturer guidelines.

**A) Antibody:** N-terminal, Abcam, cat#ab201060

*1.1Prefrontal Cortex Aβ42*

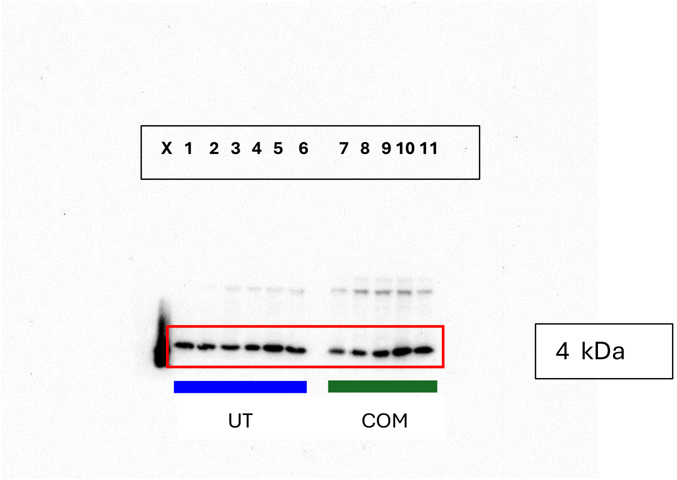


*1.2 β-Actin*

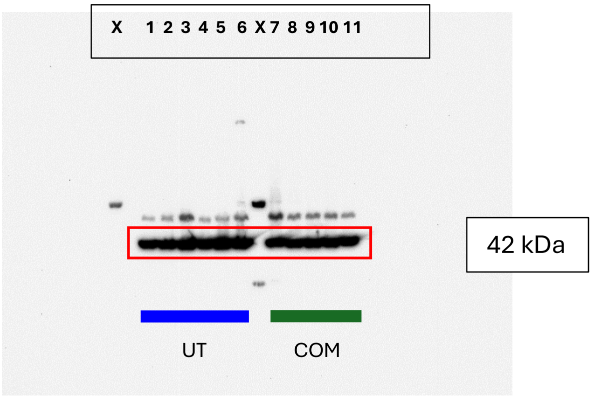

*2.1Cerebellum Aβ42*

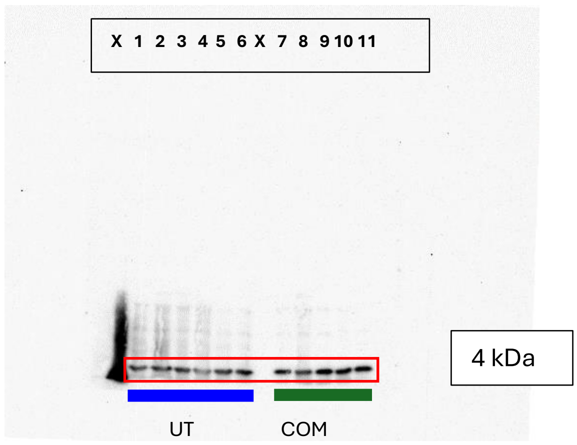


*2.2 β-Actin*

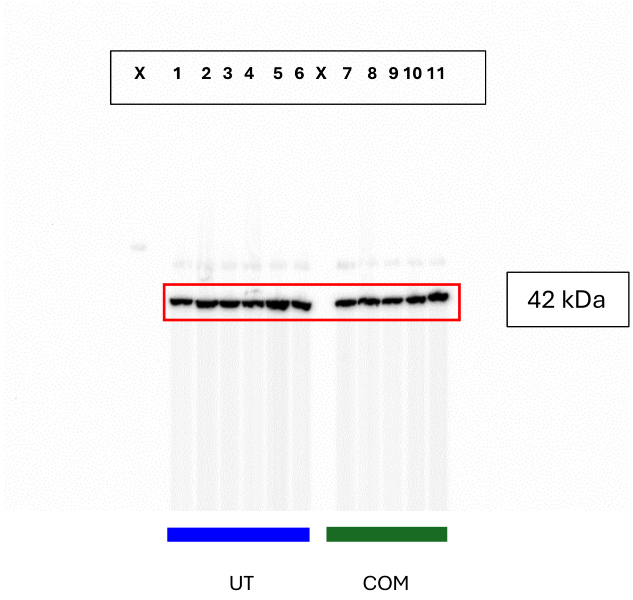

*3.1 Hippocampus Aβ42*

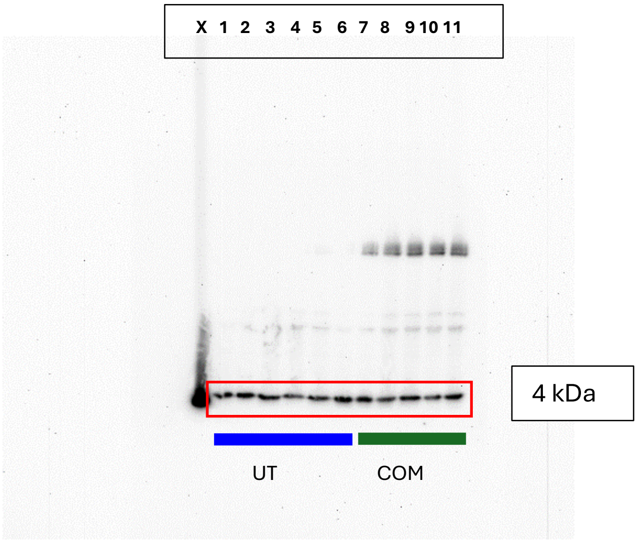


*3.2 β-Actin*

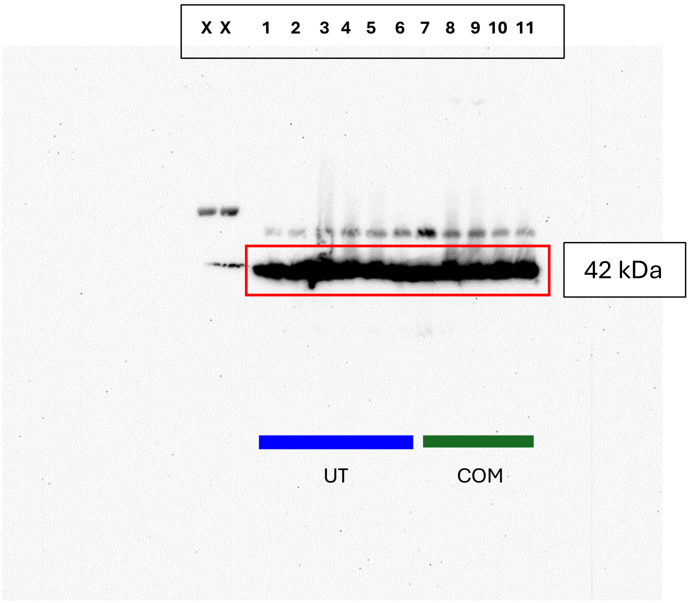

**B) Antibody:** rabbit, ThermoFisher

*1.1 Prefrontal Cortex LYVE-1*

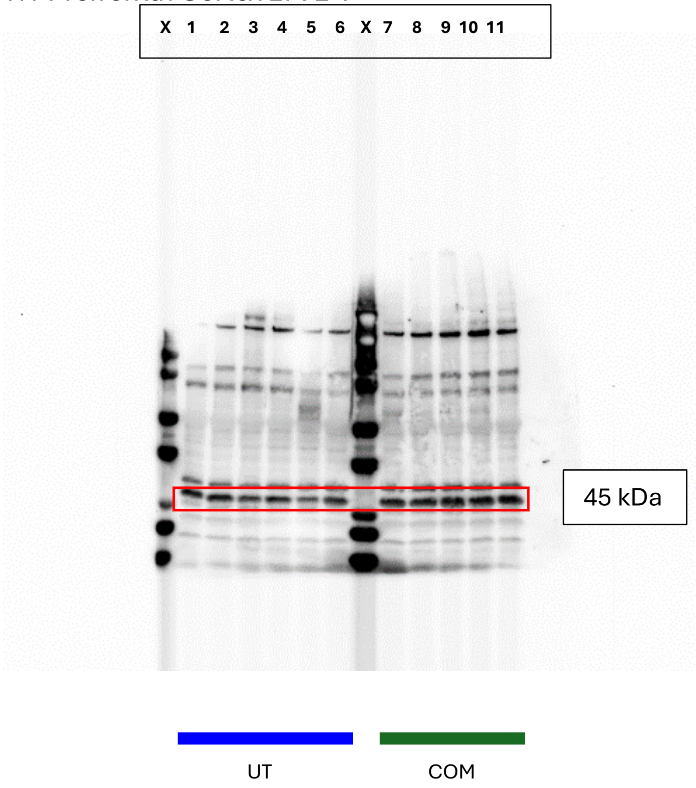

*1.2 β-Actin*

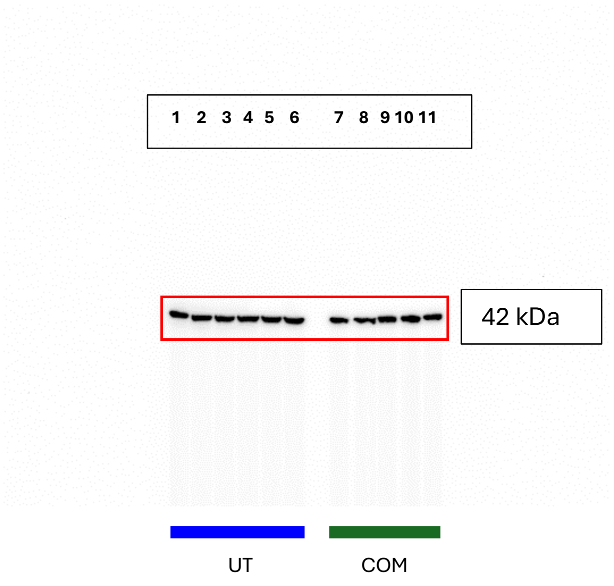


*2.1 Cerebellum LYVE-1*

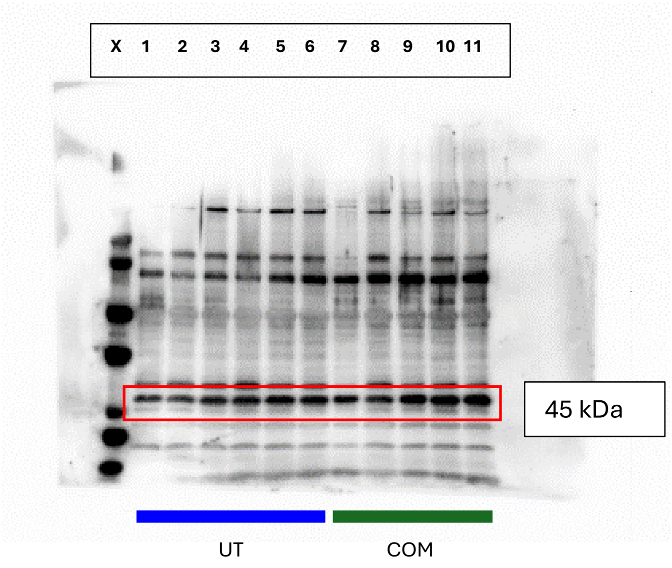

*2.2 β-Actin*

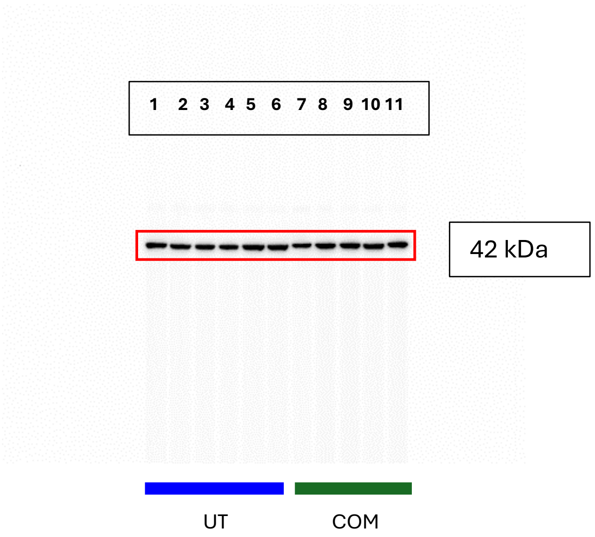


*3.1 Hippocampus LYVE-1*

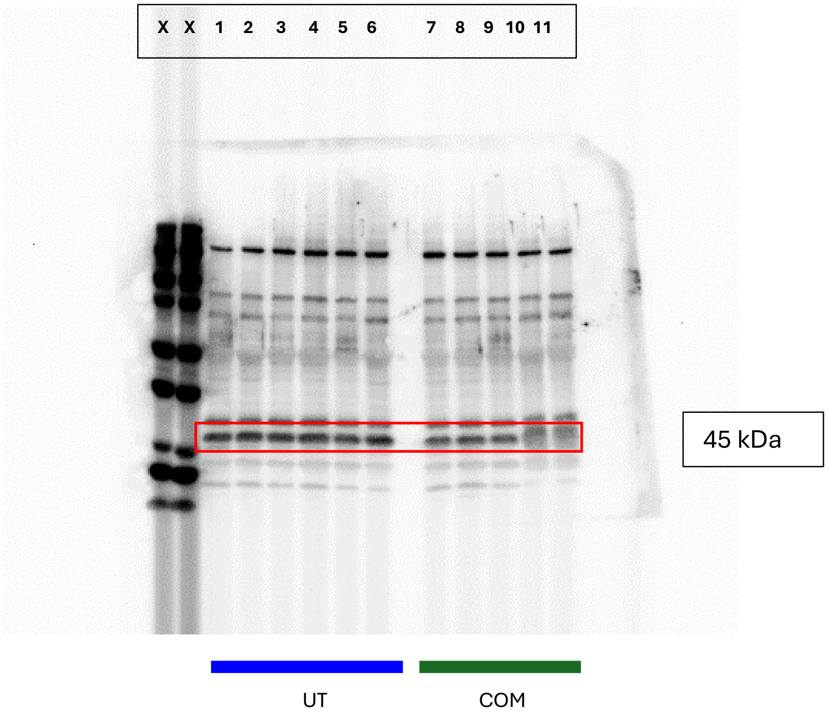

*3.2 β-Actin*

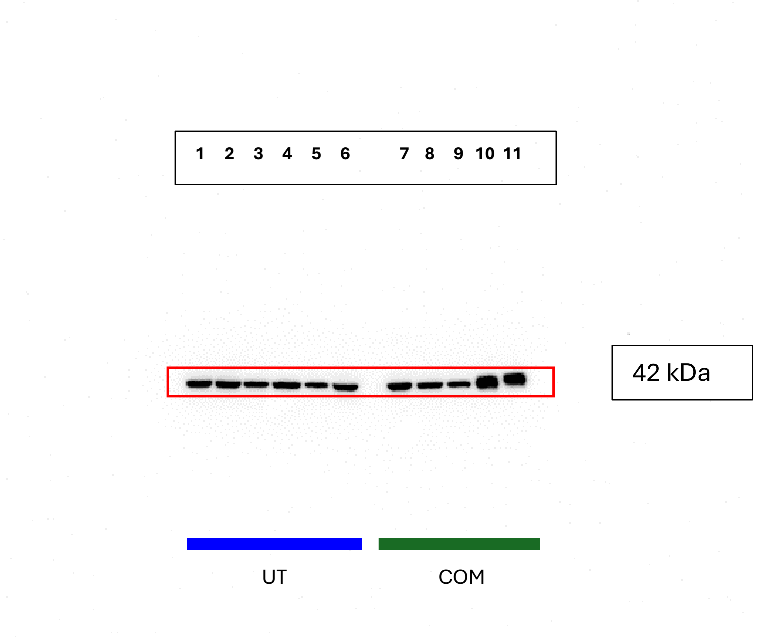


**C) Antibody:** mouse IgG2a, Millipore

*1.1 Prefrontal Cortex GAD-67*

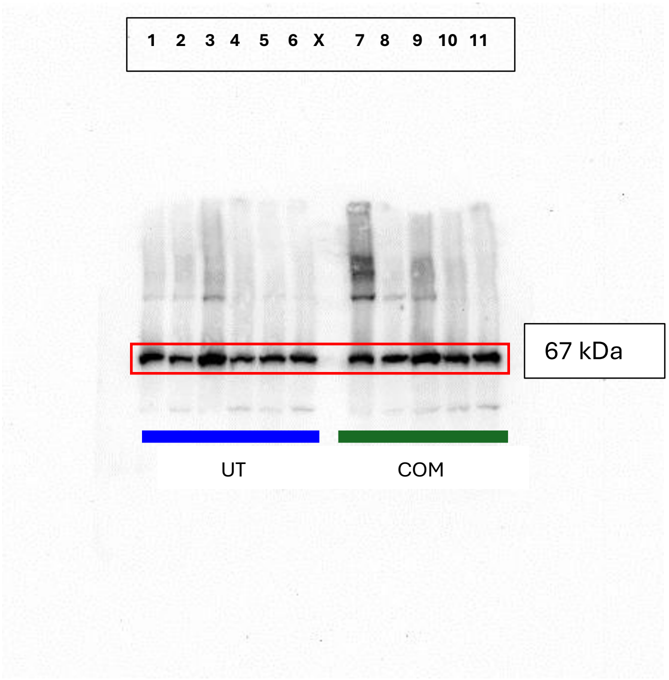

*1.2 β-Actin*

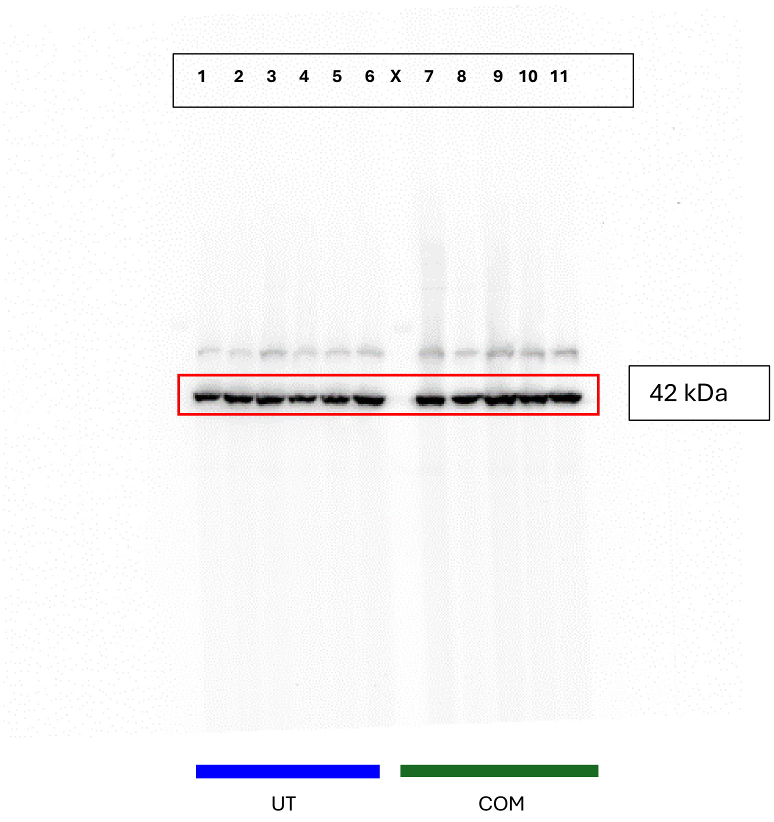


*2.1 Cerebellum GAD-67*

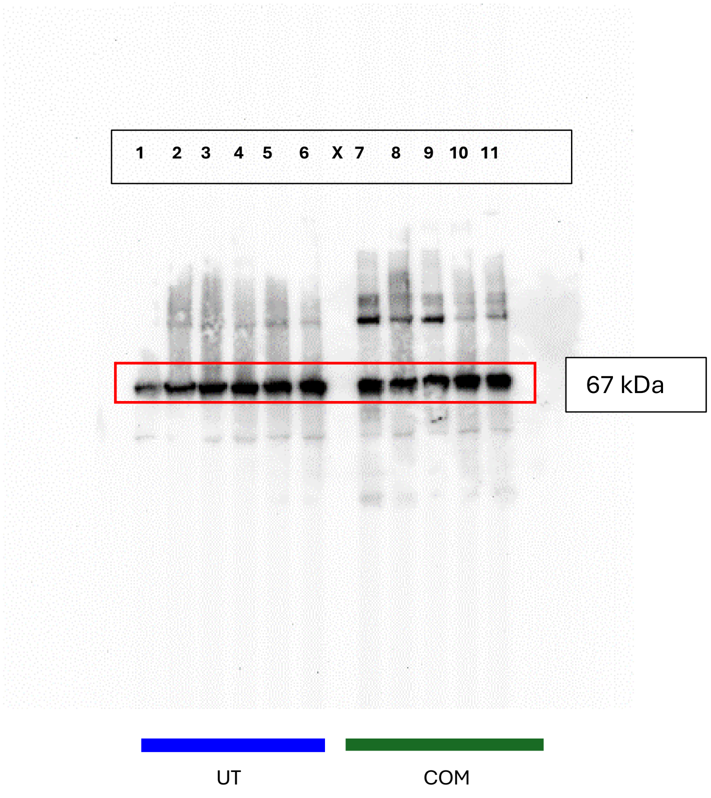

*2.2 β-Actin*

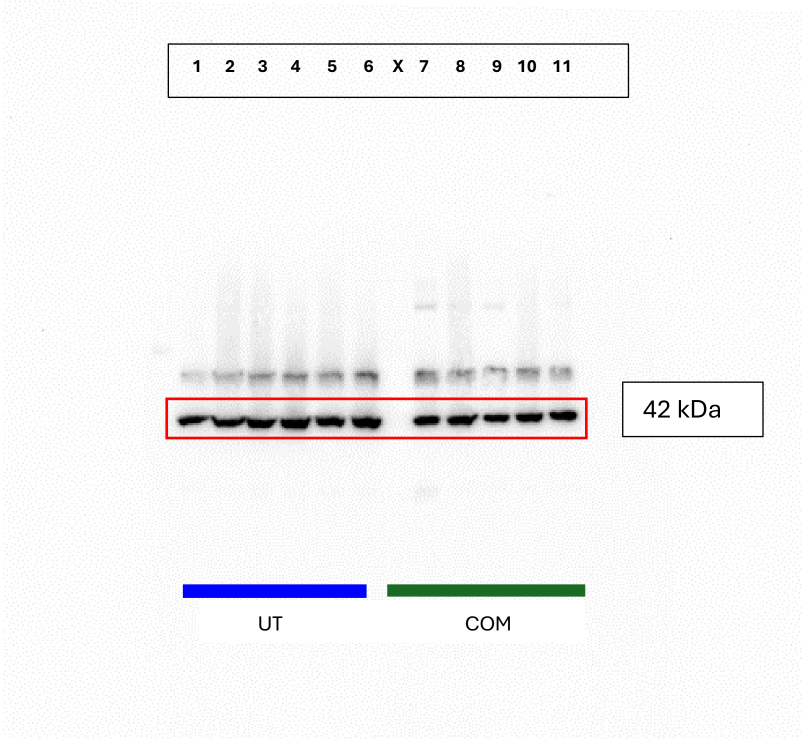


*3.1 Hippocampus GAD-67*

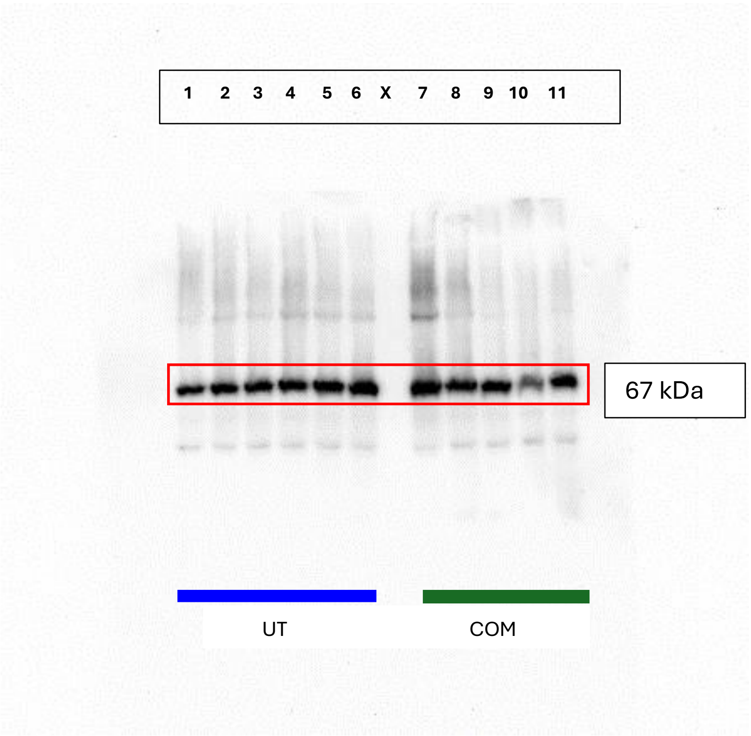

*3.2 β-Actin*

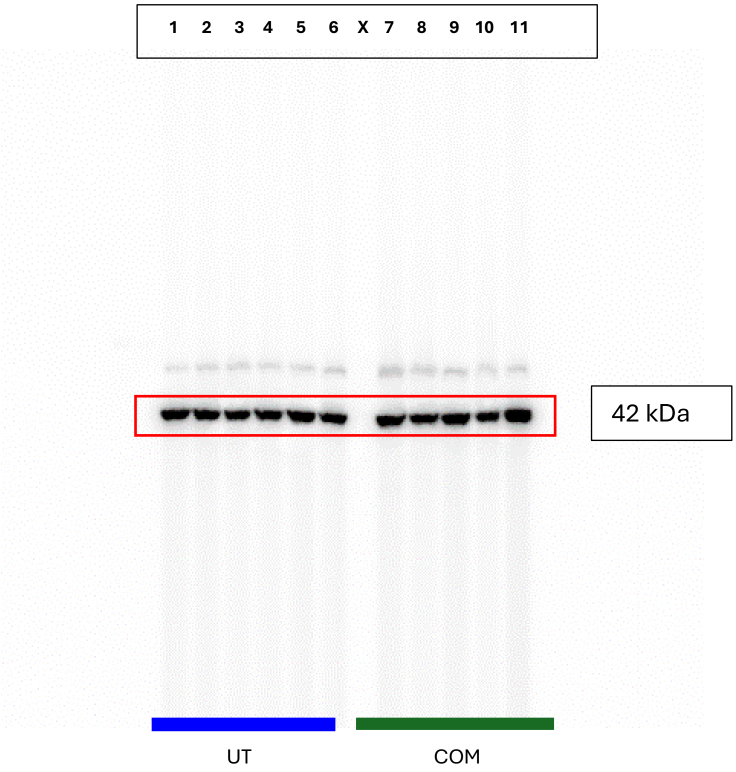

**D) Antibody: DIF8E, cell signaling**

*Prefrontal Cortex AQP4 48*

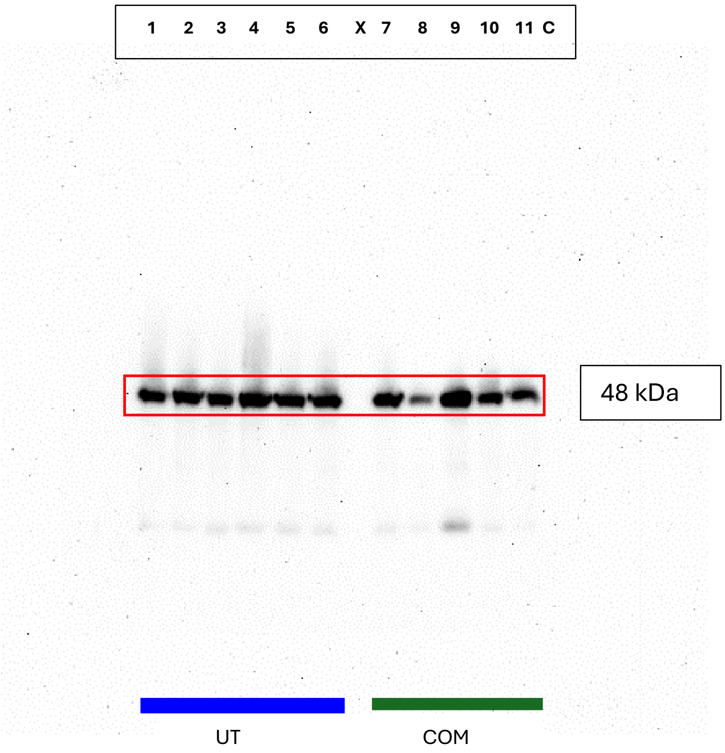


*β-Actin*

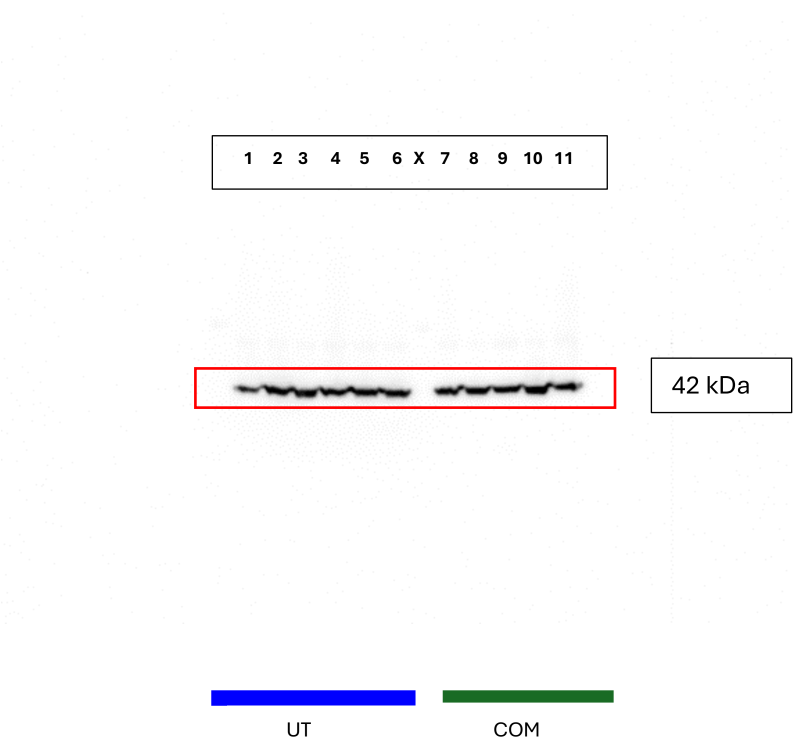
