## Supplemental document 6 for "Effect of quantified cranial osteopathic manipulation on wild type and transgenic rat models of Alzheimer’s disease"

| | | | Neurological diseases and disorders | | | | |
|---|---|---|---|---|---|---|---|
| **No.** | **Gene Symbol** | **Gene Name** | **Alzheimer's** | **Dementia** | **Epilepsy and other seizure disorders** | **Parkinson's and other movement disorders** | **Psychiatric disorders (e.g. Schizophrenia, depression, addictiion disorders, ADHD, learning disability, etc.)** |
| 1. | UBE2K | ubiquitin-conjugating enzyme E2K | x | | | | x |
| 2. | Acaca | Acetyl-CoA carboxylase 1 | X | | | | |
| 3. | CACNA1B | calcium voltage-gated channel subunit alpha1 B | | | X | | x |
| 4. | Akt | Non-specific serine/threonine protein kinase | x | | x | x | x |
| 5. | Dync1li1 | Dynein light intermediate chain | X | | | X | X |
| 6. | PAK3 | p21 (RAC1) activated kinase 3 | x | | | | X |
| 7. | Rnh1 | ribonuclease/angiogenin inhibitor 1 | | | X | | |
| 8. | VAC14 | VAC14 component of PIKFYVE complex | | | | X | |
| 9. | Tpm2 | tropomyosin 2 | | | | | X |
| 10. | Ptrh2 | peptidyl-tRNA hydrolase 2 | | | | | X |
| 11. | Syngap1 | synaptic Ras GTPase activating protein 1 | | | | | X |
| 12. | Psmc4 | proteasome 26S subunit, ATPase 4 | X | | | X | |

| | | | | | | | |
|---|---|---|---|---|---|---|---|
| 13. | Tpt1 | tumor protein, translationally-controlled 1 | | x | | | |
| 14. | Mars1 | methionyl-tRNA synthetase 1 | X | | | X | |
| 15. | THNSL1 | threonine synthase-like 2 | X | | | | |
| 16. | Krt5 | keratin 5 | X | | | | |
| 17. | Slc6a9 | solute carrier family 6 member 9 | | | x | x | |
| 18. | Srgap2 | SLIT-ROBO Rho GTPase activating protein 2 | | X | x | | |
| 19 | Mri1 | methylthioribose-1-phosphate isomerase 1 | X | | x | | |
| 20. | Faf2 | Fas associated factor family member 2 | X | | | | |
| 21. | Ctnna1 | catenin alpha 1 | X | | | | X |
| 22. | Ppip5k1 | diphosphoinositol pentakisphosphate kina se 1 | X | | | | |
| 23. | Ybx1 | Y box binding protein 1 | X | | x | x | X |
| 24. | Stxbp6 | syntaxin binding protein 6 | | | | | x |

| | | | | | | | |
|---|---|---|---|---|---|---|---|
| 25. | CTNNA2 | Alpha N-catenin | | | | | x |
| 26. | Grn | granulin precursor | x | x | | x | x |
| 27. | Myo18a | myosin XVIIIa | x | | | x | |
| 28. | Serpina3c | serine (or cysteine) proteinase inhibitor, clade A, member 3C | x | x | | x | x |
| 29. | Rps6 | ribosomal protein S6 | x | | | x | x |
| 30. | Mtmr2 | myotubularin related protein 2 [ Rattus norvegicus | | | | x | |
| 31. | Hcn2 | hyperpolarization activated cyclic nucleotide gated potassium and sodium channel 2 | | | x | | x |
| 32. | Actb | actin, beta | | | | x | x |
| 33. | Atp2c1 | ATPase secretory pathway Ca2+ transporting 1 | x | | | | |
| 34. | Nisch | nischarin | x | | | | x |
